# Gradual progression from sensory to task-related processing in cerebral cortex

**DOI:** 10.1101/195602

**Authors:** Scott L. Brincat, Markus Siegel, Constantin von Nicolai, Earl K. Miller

## Abstract

Somewhere along the cortical hierarchy, behaviorally relevant information is distilled from raw sensory inputs. We examined how this transformation progresses along multiple levels of the hierarchy by comparing neural representations in visual, temporal, parietal, and frontal cortices in monkeys categorizing across three visual domains (shape, motion direction, color). Representations in visual areas MT and V4 were tightly linked to external sensory inputs. In contrast, lateral prefrontal cortex (PFC) largely represented the abstracted behavioral relevance of stimuli (task rule, motion category, color category). Intermediate-level areas—posterior inferotemporal (PIT), lateral intraparietal (LIP), and frontal eye fields (FEF)—exhibited mixed representations. While the distribution of sensory information across areas aligned well with classical functional divisions—MT carried stronger motion information, V4 and PIT carried stronger color and shape information—categorical abstraction did not, suggesting these areas may participate in different networks for stimulus-driven and cognitive functions. Paralleling these representational differences, the dimensionality of neural population activity decreased progressively from sensory to intermediate to frontal cortex. This shows how raw sensory representations are transformed into behaviorally relevant abstractions and suggests that the dimensionality of neural activity in higher cortical regions may be specific to their current task.

**Significance statement:** The earliest stages of processing in cerebral cortex reflect a relatively faithful copy of sensory inputs, but intelligent behavior requires abstracting behaviorally relevant concepts and categories. We examined how this transformation progresses through multiple levels of the cortical hierarchy by comparing neural representations in six cortical areas in monkeys categorizing across three visual domains. We found that categorical abstraction occurred in a gradual fashion across the cortical hierarchy and reached an apex in prefrontal cortex. Categorical coding did not respect classical models of large-scale cortical organization. The dimensionality of neural population activity was reduced in parallel with these representational changes. Our results shed light on how raw sensory inputs are transformed into behaviorally relevant abstractions.

## Introduction

Neural representations at the earliest stages of cortical processing reflect a relatively faithful copy of sensory inputs, but intelligent behavior requires abstracting the behaviorally relevant elements from sensory inputs. The sensory continuum often needs to be parsed into categories, such as dividing continuous color variations of fruits into “ripe” and “unripe”. Arbitrary sensory stimuli can also be functionally associated to acquire the same meaning (e.g. the diverse stimuli grouped into the category “food”). Impaired or atypical categorization is a hallmark of disorders such as autism (1) and schizophrenia (2). Understanding its neural basis could provide pathways to early diagnosis and treatment.

Abstract categorical representations can be found in areas at or near the top of the cortical hierarchy, such as lateral prefrontal cortex (3–7), posterior parietal cortex (7–9), and the medial temporal lobe (10). Less well understood are the processing steps that transform bottom-up sensory inputs into these task-related, and thus top-down, representations. We therefore recorded from multiple regions along the cortical hierarchy in macaque monkeys performing a multidimensional categorization task. In a previous report on this dataset (11), we showed evidence that sensory signals flow in a bottom-up direction from visual to frontal cortex, while signals for the monkeys’ behavioral choice flow in a top-down direction from frontoparietal to visual cortex.

Here, we exploit the category structure of this task to investigate the degree to which visual representations in six cortical areas reflect bottom-up sensory inputs or the learned categories they are grouped into. The task required binary categorization of stimuli continuously varying along two distinct sensory domains—motion direction and color—and arbitrary grouping of a set of shape cues that signaled which feature (motion or color) should be categorized on each trial. We recorded isolated neurons simultaneously from six cortical areas (Fig. 1D) from both the dorsal and ventral visual processing streams, including frontal (lateral prefrontal cortex, PFC; frontal eye fields, FEF), parietal (lateral intraparietal area, LIP), temporal (posterior inferior temporal cortex, PIT) and visual (areas V4 and MT) cortices. Our results suggest that categorization occurs in a gradual fashion across the cortical hierarchy, reaching its apex in prefrontal cortex; that categorical coding does not always correlate with classical functional divisions; and that the dimensionality of cortical activity decreases in parallel with the reduction of continuous sensory stimuli to categorical groupings.

**Figure 1.**
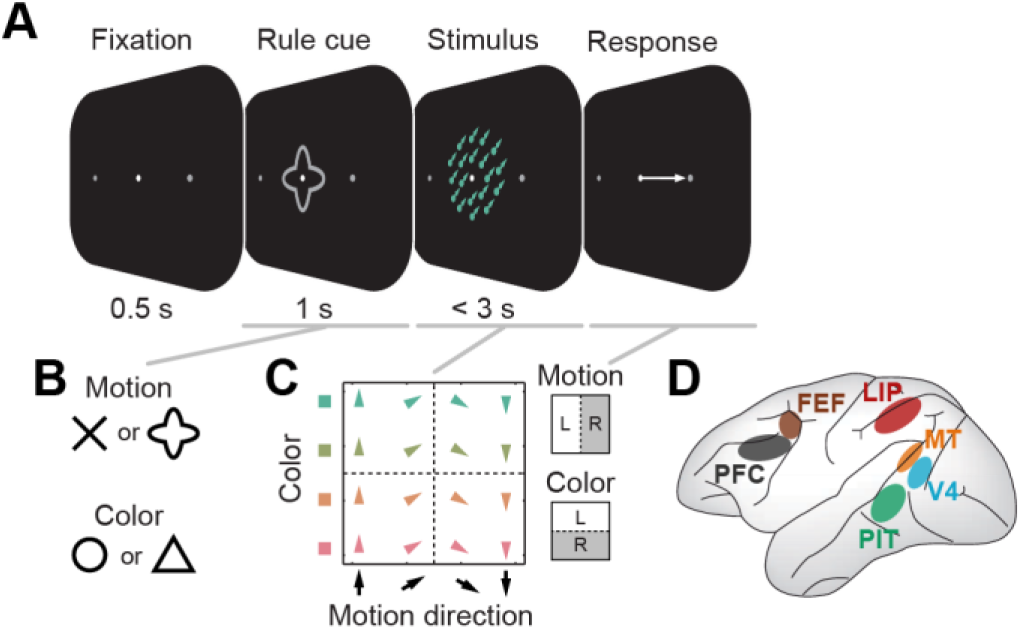
– Experimental design (*A*) Trial sequence for multidimensional visual categorization task. On each trial, the monkeys categorized either the motion direction or color of a random-dot stimulus. This stimulus was immediately preceded by a symbolic visual shape cue that instructed which feature (motion or color) to categorize for that trial. The monkey responded with a leftward or rightward saccade during the 3 s stimulus. (*B*) Either of two different cue shapes was used to instruct each task rule, in order to dissociate cue- and task-rule-related activity. (*C*) Stimuli systematically sampled motion direction (upward to downward) and color (green to red). Each color category comprised two distinct colors and each motion category comprised two distinct motion directions (additional ambiguous stimuli on the category boundaries were not analyzed here, due to our focus on categoricality). Dashed lines indicate category boundaries. For each task rule, the two categories had a fixed mapping to a leftward (L) or rightward (R) saccadic response. (*D*) Illustration of sampled brain regions: lateral prefrontal cortex (PFC), frontal eye fields (FEF), lateral intraparietal (LIP), posterior inferotemporal (PIT), V4, and MT.

## Results

Our main interest was to track the transformation of visual inputs from more sensory (bottom-up) representations to task-related (top-down) representations. On each trial of our multidimensional categorization task (Fig. 1A), a visual shape cue instructed the monkeywhether to categorize a subsequently presented colored, moving random-dot stimulus based on its color (“greenish” vs. “reddish”) or direction of motion (upward vs. downward), and report the cued category with a leftward or rightward saccade. Therefore, it probed three different types of sensory inputs: shape, motion, and color. Two of the shapes (arbitrarily chosen) cued motion categorization, while the other two cued color categorization (Fig. 1B,C). Thus, four different shapes were arbitrarily grouped into pairs by virtue of them cueing the same task rule, and four continuously varying colors and directions of motion were arbitrarily divided by a sharp boundary (Fig. 1C; additional colors/motions on category boundaries (11) were excluded from the present analyses, which require an unambiguous category assignment).

We exploited the mapping in each domain from two stimulus items (cue shapes, directions, or colors) to each categorical grouping (task rule, motion category, or color category) to dissociate stimulus-related (sensory) and task-related (categorical) effects. Purely categorical neural activity would differentiate between categories (i.e. have “preferred” responses for both items in the same category), but show no differences between items within each category. Purely sensory activity would instead differentiate between stimulus items without regard to the learned categorical divisions.

We quantified this by fitting each neuron’s spike rate, at each time point, with a linear model that partitioned across-trial rate variance into between-category and within-category effects (see “Variance-partitioning model” section in SI Methods for details). The model included three orthogonal contrast terms for each task domain (Fig. 2A). One contrast (blue) reflected the actual task-relevant of stimulus items (cue shapes, directions, or colors) into categories, and thus captured between-category variance. The other contrasts (gray) reflected the two other possible, non-task-relevant paired groupings of items, and captured all within-category variance. These three terms together capture all data variance in the given task domain. We wished to measure how much of that variance, for each domain and studied brain region, was attributable to categorical coding—its *categoricality.* Note that simply measuring the between-category variance would result in a biased estimate of categoricality—it is non-zero even for neural populations with sensory tuning for single stimulus items or for arbitrary subsets of items (Supplementary Fig. S1E,F).

**Figure 2.**
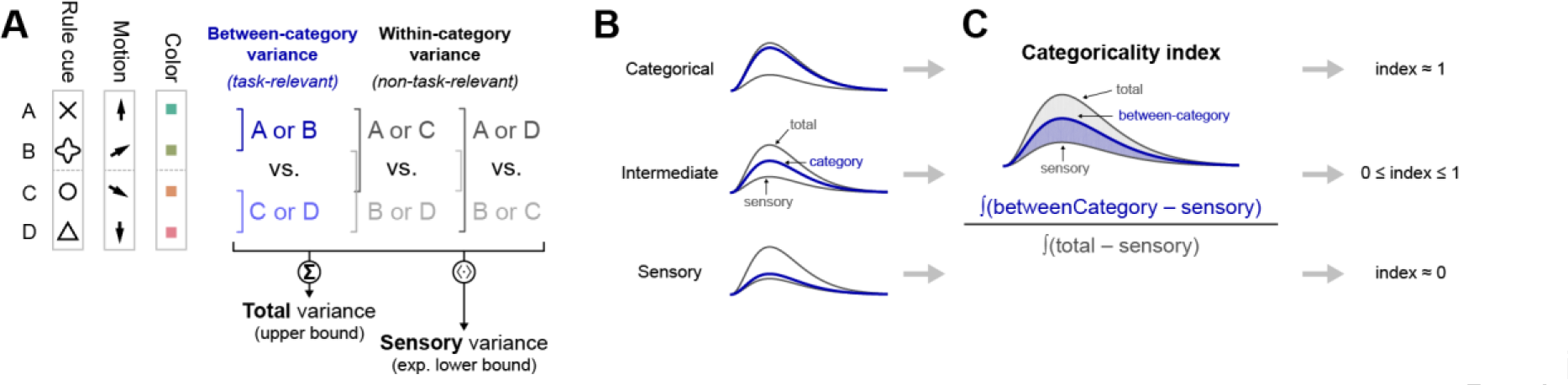
– Illustration of analysis methods (*A*) Spike rate variance for each task variable (task cues/rules, motion directions, and colors) was partitioned into three orthogonal contrasts. One contrast (blue) reflected the actual task-relevant grouping of stimulus items (cue shapes, directions, or colors) into categories, and thus captured between-category variance. The other contrasts (gray) reflected the two other possible, non-task-relevant paired groupings of items, and together captured all within-category variance. An additional term in the analysis (not depicted) partitions out variance related to the behavioral choice (left vs. right saccade). See “Variance-partitioning model” section in SI Methods for details. (*B*) The sum of variances for all three contrasts (“Σ” in panel A) bounds the between-category variance—they can be equal only for a perfectly categorical neural population with zero within-category variance (top). A purely sensory-driven population would instead have equal variance for all three contrasts, and thus between-category variance would equal the average (“〈·〉)” in panel *A*) of all three contrasts (bottom). (*C*) A categoricality index measured where actual neural populations fell between these extremes, in terms of the area between the between-category and sensory lower-bound time series, expressed as a fraction of the full area between the upper and lower bounds. Values of 0 and 1 correspond to purely sensory and purely categorical populations, respectively. See “Categoricality index” section in SI Methods for details.

Instead, we estimated where the between-category variance of each neural population fell between the predictions of purely sensory and purely categorical coding. Note that the sum of variances for all three model terms bounds the between-category variance—they can be equal only for a perfectly categorical population with zero within-category variance (Fig. 2B, top). A purely sensory-driven population would instead have equal variance for all three contrasts, and thus between-category variance would equal the average of all three terms (Fig. 2B, bottom). To measure where neural populations fall between these extremes, we computed a “categoricality index” equal to the area between the between-category and sensory (lower-bound) time series, expressed as a fraction of the full area between the total (upper-bound) and sensory (lower-bound) time series (Fig. 2C). It can be shown this is also equivalent to the between-category variance minus the average of within-category variance terms (the statistic used in our prior publication (11)), normalized by the total domain variance (see “Categoricality index” section in SI Methods for details). This index is a specific measure of how categorical a neural population is, and ranges from 1 for a perfectly categorical population to 0 for a purely sensory population. (Negative values are possible if within-category variance is greater than between-category variance, i.e. for populations that specifically reflect within-category differences.)

Within the context of each task rule, motion and color categories were—by design—inextricably linked with the monkey’s behavioral choice (e.g., under the color rule, greenish and reddish colors always mandated leftward and rightward saccades, respectively). Though this identity relationship is broken when both task rules are considered together, there remains a partial correlation between these task variables. To partition out choice effects from the category effects of interest, we also included in the fitted models a term reflecting behavioral choice. Category effects are thus measured in terms of their additional variance explained once choice effects are already accounted for (12).

To validate our analysis, we first assayed its properties on synthesized neural activity with known ground truth (see “Neural simulations” sections in SI Methods and SI Results for details). We show that our categoricality index reliably reports the relative weights of simulated sensory and categorical signals (Supplementary Fig. S1A); that it is relatively insensitive to coupled changes in both sensory and categorical signals in concert (Fig. S1B,C); and that it is relatively insensitive to simulated choice effects (Fig. S1D).

Shape information We first examined representations of the four cue shapes instructing the task rule in effect on each trial.

All sampled cortical areas—MT, V4, PIT, LIP, FEF, and PFC—conveyed significant information about the rule cues, as measured by the total spike rate variance explained by cues (Fig. 3A,B; *p* < 0.01, one-sample bootstrap test). The strongest cue shape information was in areas PIT and V4 (Fig. 3C, *p* < 0.01 for all comparisons with other areas, two-sample bootstrap test), consistent with their well-established role in shape processing (13).

**Figure 3.**
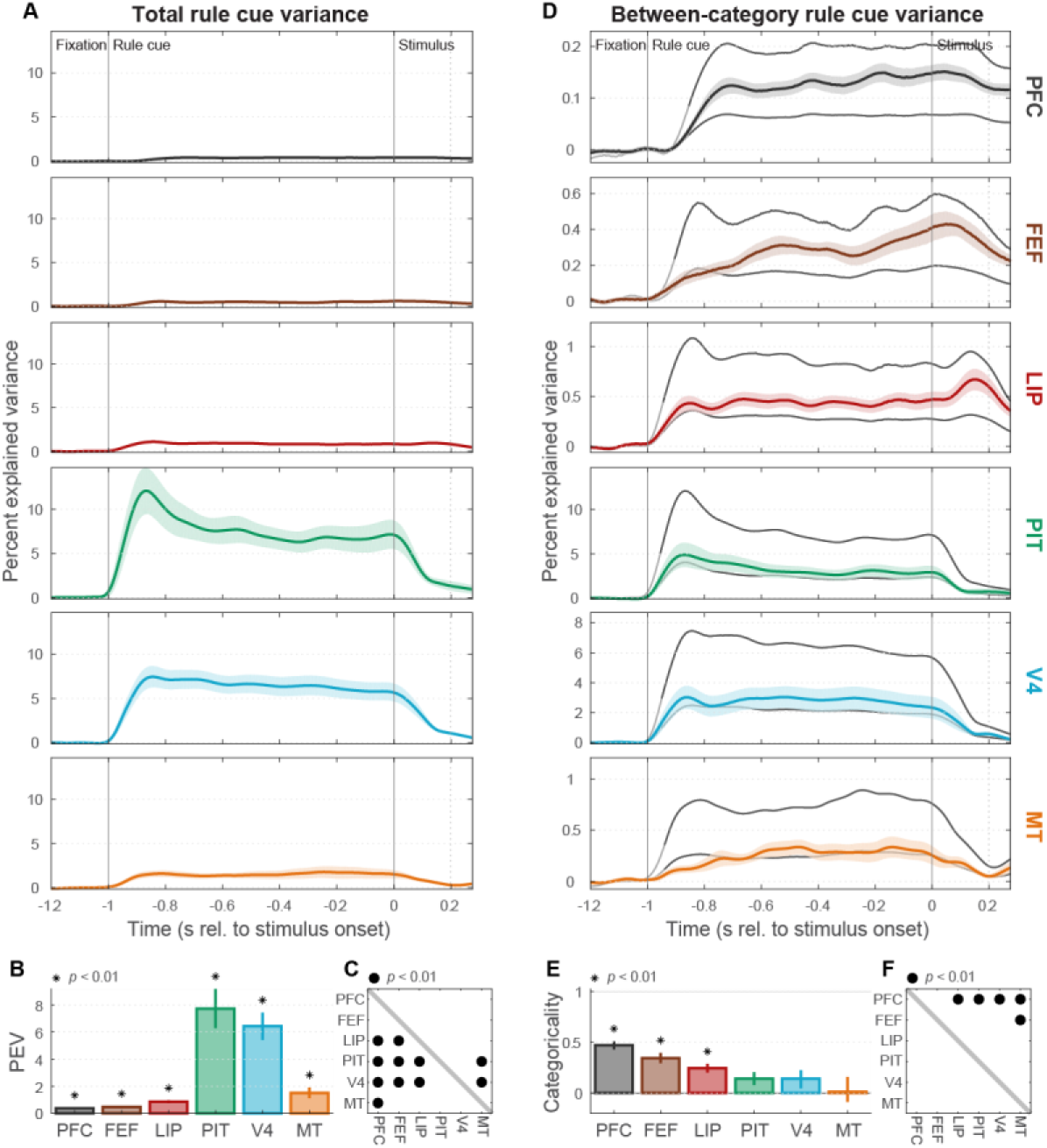
– Task rule cue representation (*A*) Population mean (± SEM) total spike rate variance explained by task rule cues (cue information), in each studied area as a function of within-trial time (referenced to the onset of the random-dot stimulus). (*B*) Summary (across-time mean ± SEM) of total rule cue variance for each area. All areas contain significant cue information (*p* < 0.01, asterisks). (*C*) Cross-area comparison matrix indicating which regions (rows) had significantly greater cue information than others (columns). Dots indicate area pairs that attained significance (*p* < 0.01). PIT and V4 contain significantly greater cue information than all other areas. (*D*) Mean (± SEM) between-category rule-cue variance (task rule information; colored curves). Gray curves indicate expected values of this statistic corresponding to a purely categorical representation of rules (upper line), and to a purely sensory representation of rule cues (lower line). The transitions from light to dark gray in these curves indicate the estimated onset latency of overall cue information, which was used as the start of the summary epoch for each area. Note differences in *y*-axis ranges from panel A. (*E*) Task rule categoricality index (± SEM) for each area, reflecting where its mean between-category rule-cue variance falls between its expected values for pure sensory (0) and categorical (1) representations. Only PFC, FEF, and LIP are significantly different from zero (*p* < 0.01, asterisks). (*F*) Indicates which regions (rows) had significantly greater task rule categoricality indices than others (columns; *p* < 0.01, dots). PFC was significantly greater than all others, except FEF.

To measure task-related (top-down) information about the task rule instructed by the cues, we partitioned out spike rate variance due to effects between task rules (“between-category variance”), and between cues instructing the same rule (“within-category variance”). Areas MT, V4, PIT all exhibited between-category variances (Fig. 3D, colored curves) that hewed closely to values expected from a pure bottom-up, sensory representation of shape (lower gray curves). We summarized these results with a “categoricality index” that measures how categorical the information conveyed by each neural population is, ranging continuously from purely sensory (0) to purely categorical (1). Task rule categoricality indices for each of these visual areas (Fig. 3E) did not differ significantly from zero (*p* > 0.01). This was true for both V4 and PIT, areas where we found strong overall cue information, as well as for MT, where there was weaker cue information. Thus, visual areas MT, V4, and PIT contained a primarily bottom-up sensory representation of the shape cues. Note that this result differs from the strong V4 and PIT task rule signals in our prior publication on this dataset (11). This is primarily due to differences in the specific questions addressed by each study, and can be reconciled by the fact that V4 and PIT do contain some task rule signals, but these signals constitute a very small fraction of the total cue variance in these areas (see “Comparison with our previous results” sections in Discussion and SI Results for details).

By contrast, PFC, FEF, and LIP all conveyed task rule information (Fig. 3D, colored curves) well above that predicted from bottom-up sensory signals (lower gray curves), and had task rule categoricality indices significantly greater than zero (Fig. 3E; *p* ≤ 1×10^−5^ for all three areas). PFC exhibited the most categorical task cue representation, significantly greater than all other areas (Fig. 3F; *p* < 0.01) except FEF (*p* = 0.05). FEF and LIP had intermediate values between PFC and the group of sensory areas (MT, V4, and PIT). All areas, including PFC, still conveyed less task rule information than expected from a purely categorical representation (Fig. 3D, upper gray curves), and had categoricality indices significantly less than one (*p* ≤ 1×10^−5^). This suggests that, unlike the visual areas, areas LIP, FEF, and particularly PFC represented the top-down meaning of the rule cues, though also retained some sensory information about them as well. Unlike the case with sensory information, where results were predictable from traditional areal divisions (shape-coding ventral stream areas PIT and V4 showed the strongest information), top-down task rule coding was observed in traditionally non-shape-coding dorsal stream areas LIP and FEF, but not in V4 or PIT.

It seems likely that some trace of the current task rule would have to persist into the stimulus period to influence how the random-dot stimulus was assigned to categories. This can be seen in the rightmost portion of Figure 3D, and we focus in on it in Supplementary Figure S2. The results indicate that cue signals in V4 and PIT decrease sharply after cue offset (Fig. S2A), and remain sensory in nature (Fig. S2D,E). In contrast, cue signals in PFC, FEF, and LIP persist relatively unabated through the stimulus period (Fig. S2D) and are in general even more categorical than during the cue period (Fig. S2E,F). These results indicate that task rule signals in frontoparietal regions (PFC, FEF, and LIP) may be involved in guiding categorical decisions about the random-dot stimuli.

### Motion direction information

Next, we turned to the random-dot stimuli that the animals had to categorize according to their direction of motion or their color. Both motion direction and color varied along a continuum, but the animals had to group them into upward/downward or greenish/reddish. Much as with the rule cues (above), we would expect bottom-up sensory signals to reflect the actual direction or color whereas task-related signals should divide them into their relevant categories. In order to discriminate signals related to stimulus category and behavioral choice, we combined data across both task rules (see above and “Variance-partitioning model” section in SI Methods for details).

First, we began with motion. Analogous to the rule cues, there were four distinct motion directions, grouped into two categories: “upward” (90° and 30°) and “downward” (−30° and −90°) categories, with rightward motion (0°) serving as the category boundary (Fig. 1C). All areas conveyed significant information about motion direction, as measured by the total spike rate variance explained by motion (Fig. 4A,B; *p* < 0.01). The strongest motion information was found in area MT (Fig. 4A,B), significantly greater than all other areas (Fig. 4C; *p* < 0.01) except PIT (*p* = 0.05), consistent with its classical role in motion processing (14).

**Figure 4.**
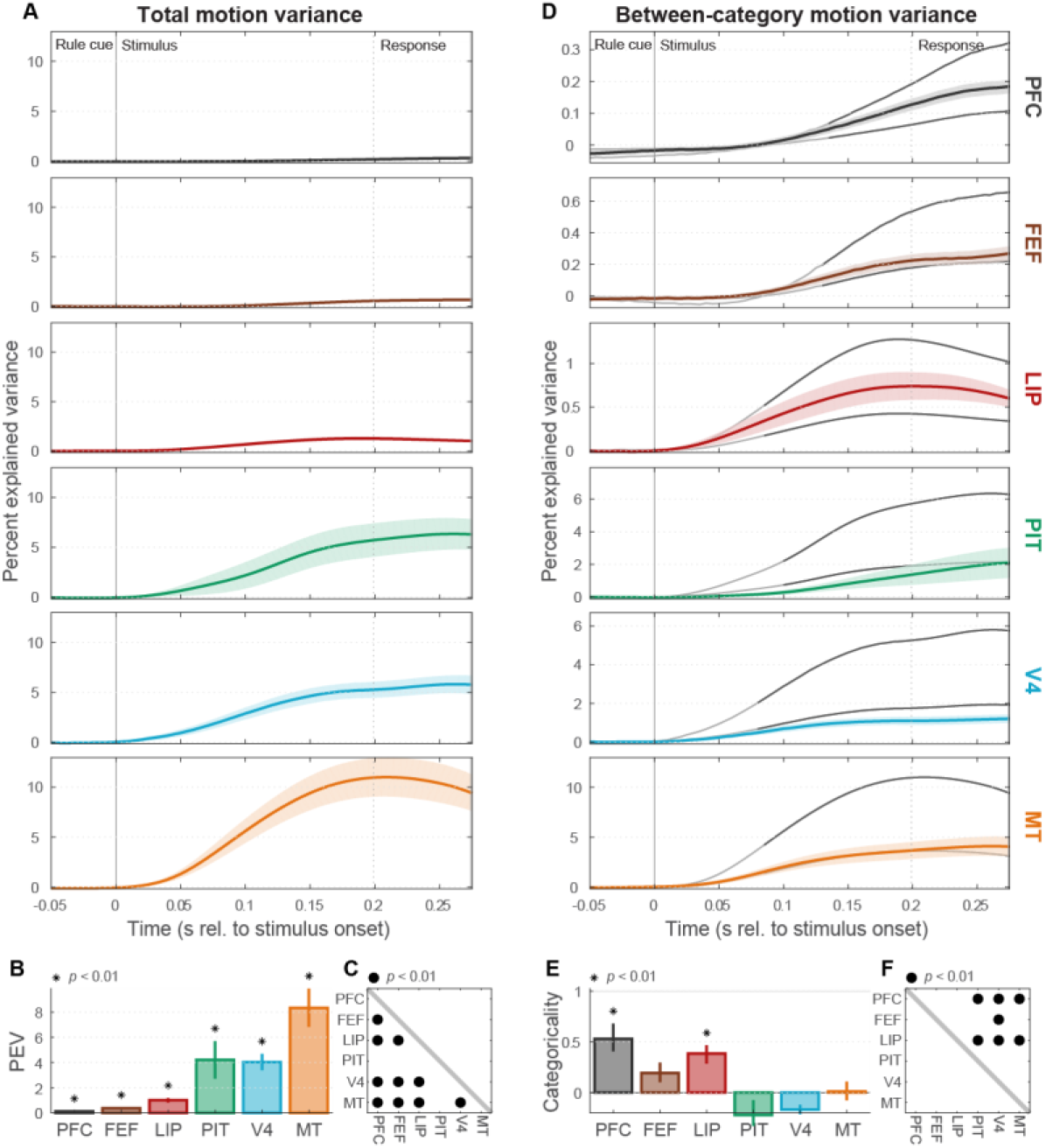
– Motion direction representation (*A*) Mean (± SEM) total rate variance explained by random-dot stimulus motion directions (motion information) in each area as a function of time (note different time axis from Fig. 3). (*B*) Summary (across-time mean ± SEM) of total motion variance for each area. All areas contain significant motion information (*p* < 0.01, asterisks), but it was strongest in MT. (*C*) Indicates which regions (rows) had significantly greater motion information than others (columns). (*D*) Mean (± SEM) between-category motion variance (motion category information). Gray curves indicate expected values for purely categorical (upper line) and purely sensory (lower line) representations of motion direction. (*E*) Motion categoricality index (± SEM) for each area, reflecting where its average between-category motion variance falls between expected values for pure sensory (0) and categorical (1) representations. Only PFC and LIP are significantly different from zero (*p* < 0.01, asterisks). (*F*) Indicates which regions (rows) had significantly greater motion categoricality indices than others (columns; *p* < 0.01, dots).

When we partitioned total motion variance into between- and within-category effects, between-category variance in MT (Fig. 4D, orange curve) closely approximated its sensory prediction (lower gray curve), and its motion categoricality index was not significantly different from zero (Fig. 4E; *p* = 0.44). The same was true of more weakly direction-selective areas V4 (*p* ≈ 1), PIT (*p* ≈ 1), and FEF (*p* = 0.03). Thus, the motion information carried by MT, V4, PIT, and FEF was largely sensory in nature.

In contrast, PFC and LIP both exhibited between-category variances (Fig. 4D, colored curves) considerably greater than their sensory predictions (lower gray curves) and motion categoricalityindices significantly greater than zero (Fig. 4E; PFC: *p* = 0.004; LIP: *p* ≤ 1×10^−5^). As with the task instruction cues, PFC showed the most categorical motion signals (Fig. 4F; significantly greater than MT, V4, and PIT, all *p* < 0.01; non-significant for FEF: *p* = 0.06 and LIP: *p* = 0.38). All areas, including PFC, remained significantly below the predictions of a purely categorical representation (Fig. 3D, upper gray curves; *p* ≤ 1×10^−5^). Thus, areas PFC and LIP conveyed top-down motion categories, but retained some sensory direction information as well. Once again, while the sensory results were consistent with traditional areal divisions (MT predictably had the strongest direction information), significant motion category information was observed in one higher-level dorsal stream area (LIP), but not in another one (FEF).

### Color information

Next, we examined neural information about the colors of the random-dot stimuli. As with motion, there were four distinct color hues grouped into two categories, “greenish” (90° and 30° hue angles) and “reddish” (−30° and −90°) with the category boundary at yellow (0°; Fig. 1C). Once again, significant color information (total color variance) could be found in all studied areas (Fig. 5A,B; *p* < 0.01). Area V4 showed the strongest color information (Fig. 5A,B), significantly greater than all other studied areas (Fig. 5C; *p* < 0.01), consistent with its established role in color processing (15). Although PIT showed the second strongest stimulus color information, it was not significantly greater than other areas (Fig. 5C; *p* > 0.01), possibly due to insufficient sampling of the relatively sparsely distributed “color patches” in and around this area (16).

**Figure 5.**
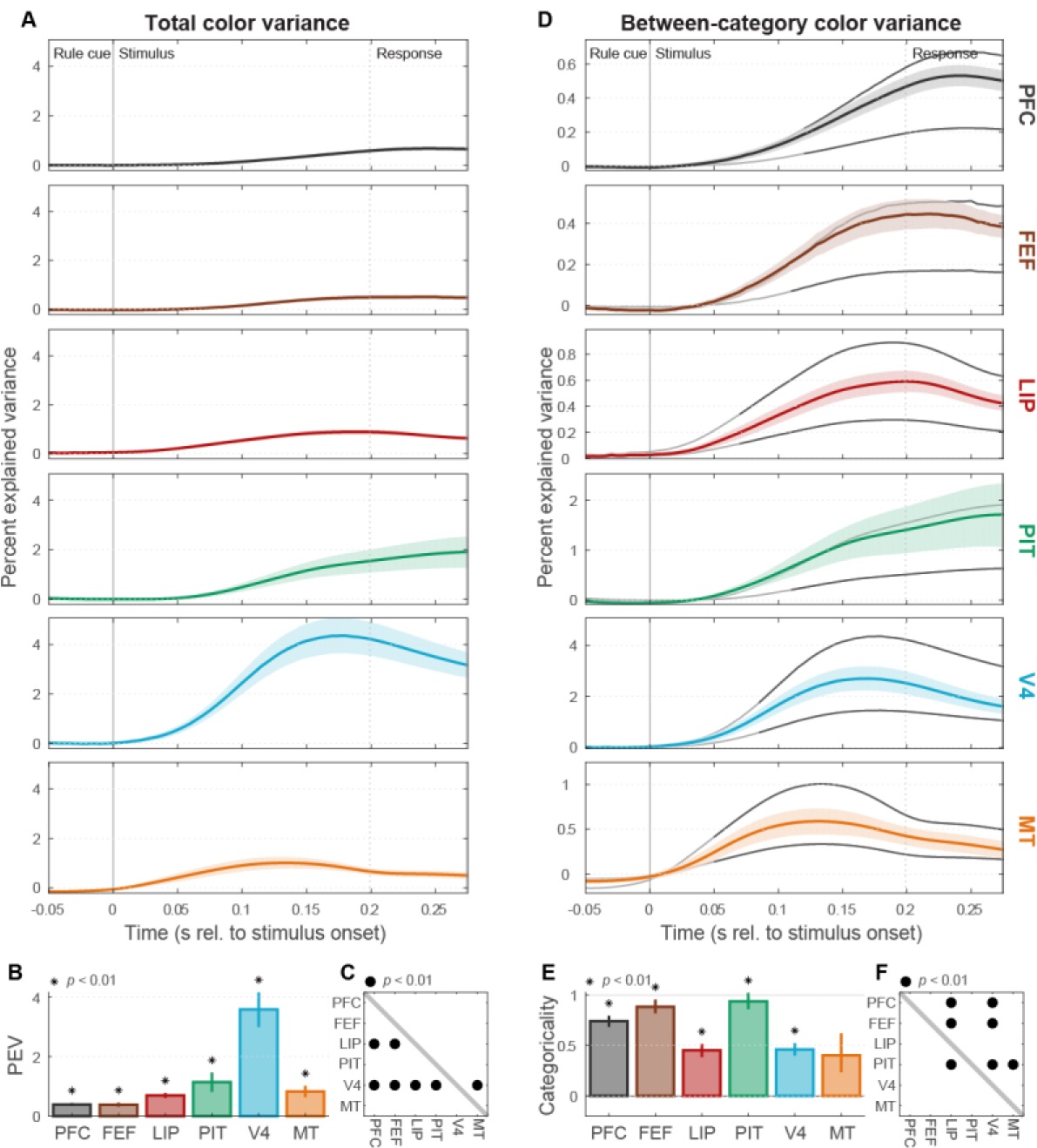
– Color representation (*A*) Mean (± SEM) total rate variance explained by random-dot stimulus colors (color information) in each area. (*B*) Summary (across-time mean ± SEM) of color information for each area. All areas contain significant information (*p* < 0.01, asterisks), but V4 carried the strongest color information. (*C*) Indicates which regions (rows) had significantly greater color information than others (columns; *p* < 0.01, dots). (*D*) Mean (± SEM) between-category color variance (color category information). Gray curves indicate expected values for purely categorical (upper line) and purely sensory (lower line) representations of color. (*E*) Color categoricality index (± SEM) for each area. All areas except MT had indices significantly greater than zero (*p* < 0.01, asterisks). (*F*) Indicates which regions (rows) had significantly greater color categoricality indices than others (columns; *p* < 0.01, dots).

Cortical representations of color were overall much more categorical than those for motion direction and rule cues, possibly due to alignment of the category structure with the red-green opponent signals arising in retinal ganglion cells and prevalent at many levels of the visual system (15). All areas except MT showed significant color categoricality indices (Fig. 5 D,E; *p* < 0.01 for all areas). PIT, FEF, and PFC all had nearly purely categorical color representations. For each of them, categorical information nearly equaled its upper bound (Fig. 5D). Their categoricality indices were significantly greater than those of LIP and V4 (Fig. 5F; *p* < 0.01), and those of PIT and FEF were not significantly different from a purely categorical representation (PIT: *p* = 0.1; FEF: *p* = 0.04). By comparison, strongly color-selective area V4, as well as weakly color-selective areas MT and LIP, were much less categorical. Thus, areas MT, V4, and LIP have a relatively bottom-up representation of color, while areas PIT, FEF, and PFC have largely categorized them into binary “greenish” and “reddish” categories. Note that, while bottom-up biases toward red-green opponent coding might have boosted the overall apparent color categoricality, it’s not obvious why such signals would be inherently stronger in higher-level areas than in V4. We also found that, once again, overall color information was fairly consistent with traditional areal divisions (V4 predictably had the strongest color information) while color categoricality exhibited mixed correlation with them: as expected, PIT was strongly categorical for color, but so was traditional dorsal stream area FEF.

### Changes in dimensionality across cortex

As a complementary assay of cortical coding properties, we examined the dimensionality of neural population activity using a noise-thresholded principal components analysis (PCA) method (17) (see “Population dimensionality analysis “ section in SI Methods for details). This analysis was previously used to measure PFC dimensionality in an object sequence memory task (18). In that context, it was found that PFC neurons contained a high-dimensional representation of task components, due to their conjunctive coding of multiple task variables (18). We asked whether high-dimensionality is an invariant property of PFC population activity, or whether it might be specific to task context. We concatenated all neurons from each studied area into a “pseudo-population” and extrapolated to larger population sizes via a condition relabeling procedure. For each area and population size, we computed the mean spike rate for each of 64 task conditions (4 rule cues × 4 motion directions × 4 colors) within the random-dot stimulus epoch, when all conditions were differentiated. The dimensionality of the space spanned by each resulting set of 64 neural population activity vectors was quantified as the number of principal components (eigenvalues) significantly greater than those estimated to be due solely to noise. As expected, estimated dimensionality grew with the size of the neural population examined but generally approximated an asymptotic value (Fig. 6A) that can be taken as an estimate of the dimensionality of the underlying (much larger) neural population. Clear differences in the estimated dimensionality were observed across areas, summarized in Fig. 6B. The highest dimensionality was observed in visual areas V4, PIT, and MT, presumably reflecting a large diversity of sensory tuning curves in these visual areas. These were followed by intermediate-level visual areas LIP and FEF. The lowest dimensionality was observed in PFC. The observed high dimensionality of area PIT was likely due in part to the inclusion of two task variables that it carried relatively strong information about: rule cues (shape) and color. When the same analysis was performed in a 16-dimensional space consisting only of 4 directions × 4 colors (Fig. 6C,D), PIT dimensionality was greatly reduced and was similar to that of LIP.

**Figure 6.**
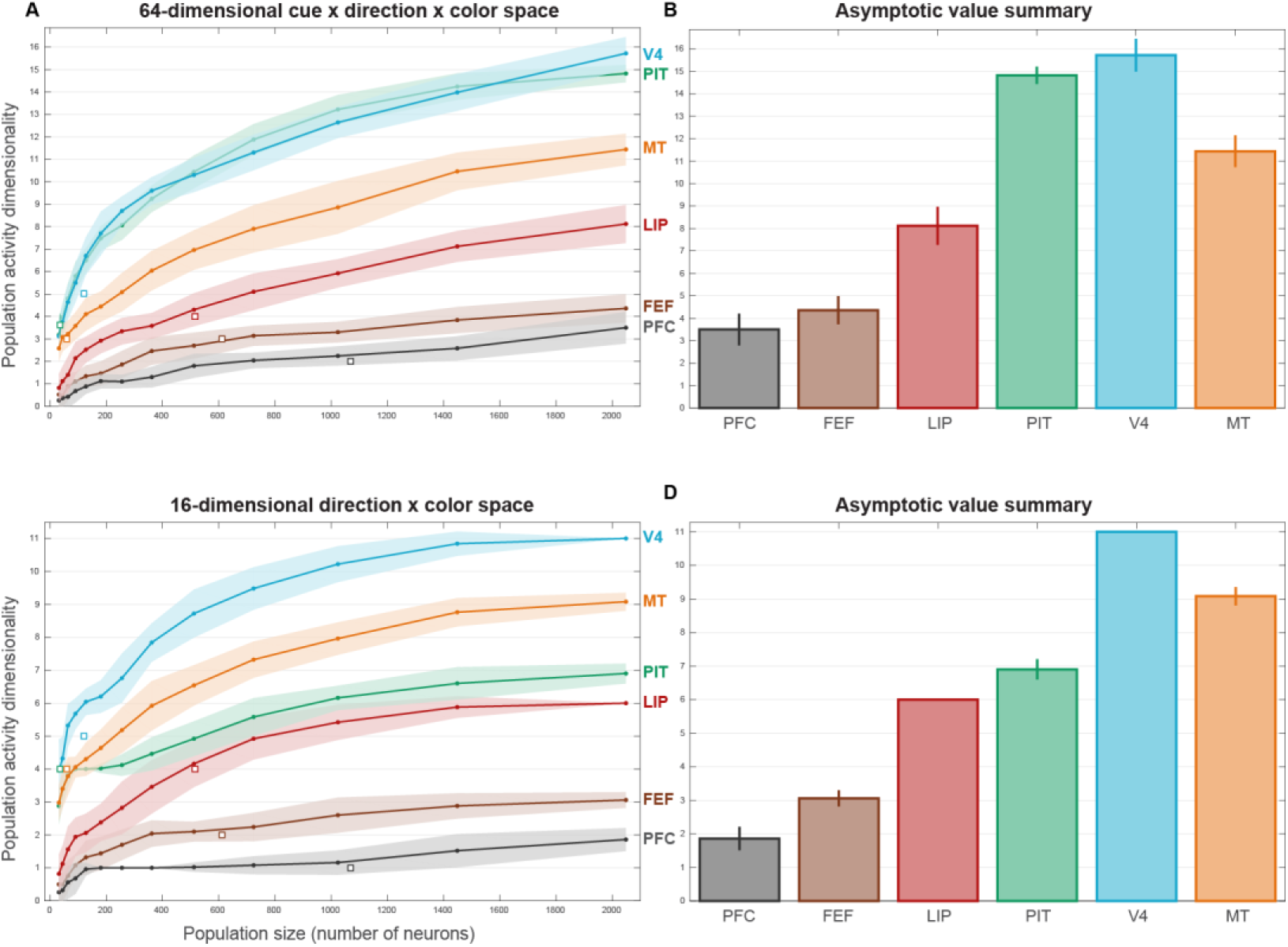
– Population activity dimensionality (*A*) Dimensionality (mean ± SEM) of neural population activity as a function of extrapolated population size, for each studied area. Dimensionality was estimated by noise-thresholded PCA within a 64-dimensional rule cue × motion direction × color space (see “Population dimensionality analysis” section in SI Methods for details). Values for the actual recorded neural populations (white squares ± SEM) were largely consistent those from the extrapolated populations. (*B*) Summary of asymptotic dimensionality values (± SEM) in 64-d space. (*C*) Dimensionality (mean ± SEM) of population activity as a function of population size, for each studied area. Dimensionality was estimated within a reduced 16-dimensional motion direction × color space (*D*) Summary of asymptotic dimensionality values (± SEM) in 16-d space. V4 and MT have the highest dimensionality, followed by PIT and LIP, and then by FEF and PFC.

Thus, the dimensionality of population activity decreased progressively up the cortical hierarchy in parallel with the gradual shift from sensory to categorical representations. Further, the estimated PFC dimensionality is close to the value that would be expected of a purely categorical representation with binary responses for each task variable (~3 for the 3-variable analysis, Fig. 6B; ~2 for the 2-variable analysis, Fig. 6D). These results suggest that high dimensionality is not an invariant property of PFC activity, but may be specific to current behavioral demands, in this case to reduce high dimensional sensory stimuli to binary categories.

## Discussion

### Abstraction of sensory inputs occurs progressively through the cortical hierarchy

Our results, summarized in Figure 7, demonstrate a gradual progression from bottom-up sensory inputs to abstracted, top-down behaviorally relevant signals as the cortical hierarchy is ascended. Across three visual domains—shape, motion direction, and color—lower-level visual areas MT and V4 conveyed strong information about sensory stimuli within their preferred domains (Fig. 7A) but showed little evidence for any abstraction beyond the raw sensory inputs (Fig. 7B). In contrast, higher-level area PFC, despite containing relatively weak information overall (Fig. 7A), showed strongly abstracted, task-relevant coding across all domains (Fig. 7B). In between, intermediate-level visual areas PIT, LIP, and FEF showed mixed representations with partially categorical coding in some domains, but not others. These results support models of cortical processing where representational transformations happen gradually across multiple cortical processing steps (19, 20), rather than in a discrete, all-or-nothing fashion.

**Figure 7.**
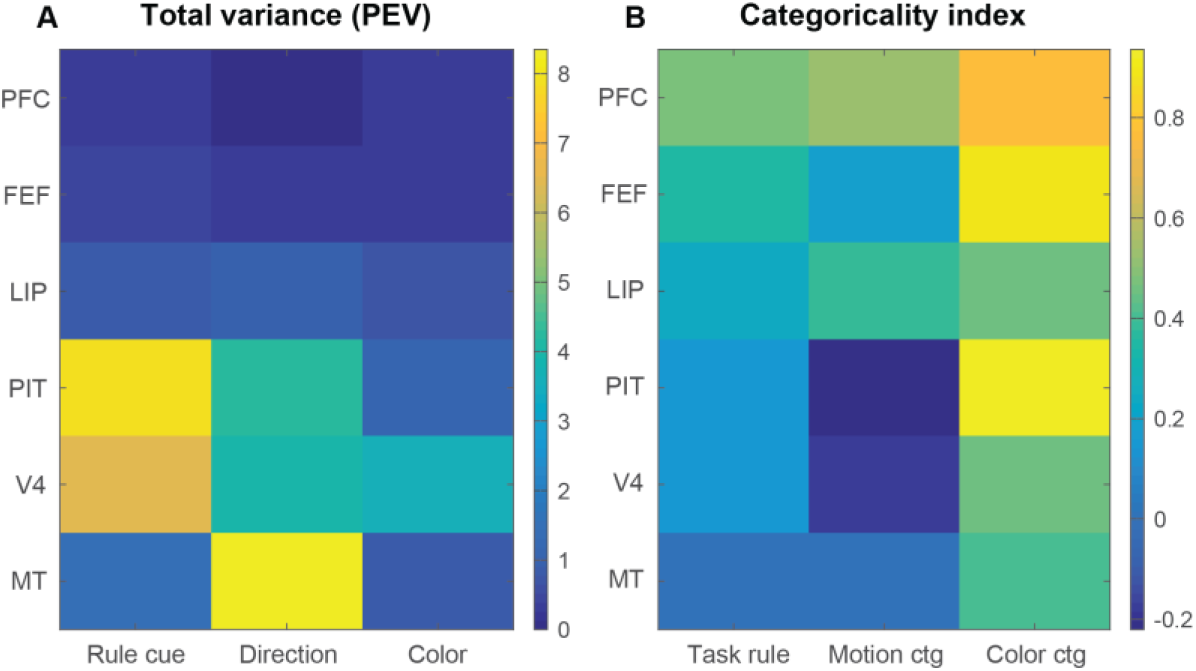
– Summary of results (*A*) Mean total variance in each studied area explained by rule cues, motions directions, and colors. (*B*) Categoricality indices for each studied area for task rules, motion categories, and color categories.

In all but a few cases (PIT and FEF for color), cortical representations—even in high-level areas—remained significantly less categorical than predicted for a purely categorical neural population. The distribution of categoricality index values across neurons in the studied populations (Supplementary Fig. S3) suggests two reasons for this. First, even mostly categorical populations contained some residual sensory-coding neurons (index values ≈ 0). Second, all studied populations contained some individual neurons whose activity differentiates both between categories and between items within each category (0 ≤ index ≤ 1). Thus, the intermediate categorical coding we observed in most areas reflected a mixture of sensory and categorical effects at both the level of single neurons and the neural population. Purely categorical signals might exist in other brain regions, such as the medial temporal lobe (10). However, some multiplexing of categorical and residual sensory signals could also have a functional role, such as permitting behavior to be driven flexibly by multiple levels of abstraction.

Note that the exact values of categoricality indices may be influenced by the particular stimuli used and are thus to some degree specific to this task. For example, the apparent strong categoricality of color representations may have been due to alignment of the category structure with the red-green opponent signals arising in retinal ganglion cells (15). However, it’s not obvious why any such bottom-up stimulus biases would be inherently stronger in higher-level areas, without being driven by a learned category structure. Thus, while we do not put strong interpretational value on the exact index values for each area and task variable, we believe their relative values accurately portray a progression from sensory-dominated to categorical coding through the cortical hierarchy.

### Comparison with our previous results

One result may appear to be somewhat at odds with our prior publication on this dataset (11). We previously claimed that V4 and PIT had strong task rule information (Fig. 2C in (11)). Here, we claim task rule categoricality is weak and non-significant in these areas (Fig. 3D,E). This difference lies primarily in the specific questions addressed by each study. Our previous study addressed the overall task rule information conveyed by each neural population. It therefore used a statistic that measured a debiased version of between-category variance for task cues. This measure could be high for populations conveying strong information about a domain— such as the representations of rule cue shapes in V4 and PIT— even if only a very small fraction of that information is categorical (see simulations in Fig. S1H). Here, we instead addressed how *categorical* neural representations are. We used a statistic that normalizes out overall information to measure categoricality *per se* (Fig. S1B,C). Thus, we can reconcile results from the two studies by concluding that V4 and PIT contain strong information about task cues but only a small fraction of that information is categorical. In contrast, despite the weaker overall task cue information in PFC and FEF, a substantial fraction of that information reflects the learned task rule categories. This definition accords well with both intuitive notions of categoricality and those previously proposed (3, 10). As elaborated below, it is further supported by its tight correspondence to the anatomically-defined cortical hierarchy.

### Graded cortical functional specialization

There is a longstanding debate about the degree to which the functions of different cortical regions are specialized or overlapping. The cortex’s broad interconnections and remarkable resistance to localized damage argue for more distributed, overlapping representations (21), while many studies find evidence for seemingly circumscribed functions in at least some cortical regions (22, 23). We find evidence supporting both points of view. Information about task variables was not distributed homogeneously across cortical regions. For each variable, one or two areas clearly conveyed much stronger information (Fig. 7A) than others. These results were largely predictable from the classical functions of visual cortical areas. Area MT was dominant for motion direction, consistent with its well-established role in motion processing (14). V4 was dominant for color, consistent with many reports of its robust color selectivity (15). PIT and V4 both showed strong sensory information about rule cue shapes consistent with their well-established role in shape processing (13). Thus, these results support the idea of specialized cortical representations and reconfirm some of the classical functional divisions between ventral stream and dorsal stream areas using an experimental paradigm where multiple areas from both streams were tested simultaneously across multiple stimulus domains.

On the other hand, significant information about all examined experimental variables was found in all sampled areas, supporting the idea of broadly distributed cortical representations. The fact that color and shape information can be found in dorsal stream areas MT, LIP, and FEF and motion information can be found in ventral stream areas V4 and PIT argues against any absolute functional dichotomies between cortical processing streams, consistent with previous reports (24–26). We believe this body of results supports cortical models with graded functional specialization, where cortical areas have clear innate or learned biases to represent certain attributes but retain coarse, distributed information about non-specialized attributes (27, 28).

While the overall strength of task-related information accorded well with classical divisions, the degree of top-down categorical abstraction painted a somewhat different picture. Dorsal stream area FEF exhibited a strongly categorical representation of color and task rule (derived from cue shape) but a non-categorical, sensory representation of motion direction. LIP was predictably categorical for motion but also showed a moderately categorical representation for task rule (shape) and color. While areas V4 and PIT were somewhat more predictable—they were relatively categorical for color, but not at all for motion direction—they unexpectedly exhibited little to no categorical coding for the cue-shape-derived task rule.

We quantified these observations by explaining the summary data in Figure 7 with two predictors related to large-scale cortical organization: 1) the anatomically-derived hierarchical level of each area (29), and 2) the expected functional congruence of each combination of task variable and area—positive for those consistent with classical dorsal/ventral processing stream divisions (e.g. MT and motion, V4 and color), and negative for inconsistent combinations (e.g. MT and color, V4 and motion; see “Cortical organization analysis” section in SI Methods for details). We found that both the decrease in sensory information and the increase in categorical coding across cortical areas were well explained by their anatomical hierarchical level (*p* ≤ 1×10^−5^ for both) with only a marginally significant difference between them (*p* = 0.04). In contrast, only sensory information was also significantly explained by classical processing stream divisions (*p* ≤ 1×10^−5^), while categoricality index values were not (*p* = 0.11), with a significant difference between them (*p* = 0.003). Thus, while our sensory information results confirm classical areal divisions, the degree of categorical coding is not well explained by them. These results suggest cortical regions may form different functional networks for bottom-up vs. top-down functions, putatively reflecting primarily feedforward and feedback/recurrent circuits, respectively.

### Categorization reached its apex in prefrontal cortex

Many studies have now reported neural correlates of categories in PFC (3, 5–7), and in area LIP and related areas of posterior parietal cortex (6, 7, 9). A recent study suggested that LIP might contain a stronger representation of categories that could drive categorical processing in PFC (6). We found that, for all examined domains, PFC exhibited a degree of categorical abstraction either greater than all other studied areas (task rule and motion) or not significantly different from the other most categorical area (color). For all domains, the prefrontal representation was more categorical than LIP, though this difference was significant only for task rule and color, not motion direction. On the other hand, despite being less categorical than PFC, LIP did also exhibit a significantly categorical representation for all tested domains, which was not the case for any other studied area besides PFC. We interpret these results to mean that PFC does play an important—perhaps the most important—role in categorization (see also (7)). However, LIP clearly also plays a central role, and categorization likely involves reciprocal interactions between these areas, as well as others (30, 31).

### Cortical dimensionality may be task-specific

We found a progression from high-dimensional population activity in the visual areas (V4 and MT) to low-dimensional populations in the frontal areas (PFC and FEF), paralleling the change in categoricality. We interpret this to reflect a shift from a large diversity of sensory tuning curves in visual cortex to nearly binary categorical responses in PFC.

At first blush, however, these results might seem at odds with a recent report showing prefrontal population activity is high dimensional (18). That study found that PFC neurons tend to exhibit “nonlinear mixed selectivity” for specific conjunctions of task variables and consequently PFC population activity had a dimensionality near the theoretical maximum (24 dimensions) for the studied task. However, that study employed a task involving encoding and maintenance in working memory of a sequence of visual objects and responding either via a recollection or recall probe (32). Thus, correct performance required remembering which of 12 different sequences was shown and which of two modes of behavioral output was mandated. By contrast, the task used here emphasized dimensionality *reduction.* First, four visual cues were grouped into either of two task instructions. Next, 16 random-dot stimuli (4 colors × 4 directions) was mapped onto binary color or motion categories, depending on the currently instructed task rule. Finally, the deduced category was translated into a binary response. Thus, this task, unlike the previous one, emphasized reduction of high-dimensional sensory inputs to lower-dimensional abstractions. Our results therefore suggest the possibility that prefrontal dimensionality may flexibly reflect current cognitive demands (33). Inputs may be expanded to higher dimensions when decisions depend on multiple variables but reduced to lower dimensionality when categorical abstraction is required. Thus, PFC dimensionality, like other PFC coding properties (34), appears to flexibly adapt to behavioral needs.

## Methods

Experimental methods are briefly reviewed here, but further details can be found in SI Methods, as well as in our prior publication from this dataset (11). All procedures followed the guidelines of the Massachusetts Institute of Technology Committee on Animal Care and the National Institutes of Health.

### Electrophysiological data collection

In each of 47 experimental sessions, neuronal activity was recorded simultaneously from up to 108 electrodes acutely inserted daily into up to six cortical regions (Fig. 1D): MT, V4, PIT, LIP, FEF, and lateral PFC. All analyses were based on 2414 well-isolated single neurons (MT: 60, V4: 121, PIT: 36, LIP: 516, FEF: 612, and PFC: 1069). The basic analysis was also repeated using multi-unit signals (pooling together all threshold-crossing spikes on each electrode), with very similar results (Supplementary Fig. S4). To minimize any sampling bias of neural activity, we did not prescreen neurons for responsiveness or selectivity. See “Electrophysiological data collection” section in SI Methods for details.

### Behavioral paradigm

Two adult rhesus macaques *(Macaca mulatta)* were trained to perform a multidimensional categorization task. On each trial (Fig. 1A), a visual cue instructed the monkey to perform one of two tasks: color categorization (“greenish” vs. “reddish”) or motion categorization (“upward” vs. “downward”) of a subsequently presented colored, moving random-dot stimulus. They responded via a saccade towards a target to the left (greenish/upward) or right (reddish/downward). See “Behavioral paradigm” section in SI Methods for details.

### Data analysis

#### General

For most analyses, spike trains were converted into smoothed rates (spike densities). To summarize results, we pooled rates or other derived statistics within empirically-defined epochs of interest for each task variable and area (see “General” section in SI Methods for details). Only correctly performed trials were included in analysis. All hypothesis tests used distribution-free bootstrap methods, unless otherwise noted.

#### Categoricality analysis

Our primary interest was to characterize each cortical region’s categoricality—the degree to which it reflected the raw sensory stimuli or their abstracted meaning (task rule or motion/color category). We quantified this by fitting each neuron’s spike rate, at each time point, with a linear model that partitioned across-trial rate variance within each task domain into between-category and within-category effects (Fig. 2A). We then computed a categoricality index reflecting where the observed between-category variance for each population fell between the predictions of purely sensory and categorical coding (Fig. 2B). Because overall variance within each task domain is effectively normalized out of this index, it reflects a pure measure of the categorical quality of a neural representation, similar to previous measures of category selectivity (3), but taking the reliability of neural coding into account, as it’s based on explained variance, rather than raw spike rates. See “Variance-partitioning model” and “Categoricality index” sections in SI Methods for details. Our analysis methods were validated with extensive simulations (Supplementary Fig.S1), and supported by a separate analysis comparing predictions of category and choice coding (Supplementary Fig. S5). See “Neural simulations” and “Category/choice consistency analysis” sections in SI Methods and SI Results for details.

We also measured categoricality by comparing mean spike rates for preferred categories and stimulus items within categories. The relative difference in spike rates between and within categories was generally consistent with our presented results (Supplementary Fig. S6). See “Preferred condition analysis” sections in SI Methods and SI Results for details.

#### Population dimensionality analysis

To measure the dimensionality of population activity, we estimated the number of principal components required to describe the space spanned by condition-mean neural population activity vectors (17, 18). Epoch spike rates were computed for each trial and neuron, averaged across all trials of each condition, and concatenated across all neurons and sessions into a set of neural “pseudo-population” vectors for each studied area. Dimensionality was computed as the number of principal components (eigenvalues) of each resulting matrix significantly greater than the estimated distribution of principal components due to noise. See “Population dimensionality analysis” section in SI Methods for details.

#### Cortical organization analysis

To relate our results to classical models of cortical organization, we fit the data in each of the population summary matrices of Figure 7 with a two-predictor linear model: (1) the Felleman and Van Essen (29) hierarchical level of each area, and(2) the expected functional congruence of each combination of task variable and area based on classical functional divisions—positive for consistent combinations (e.g. MT and motion), and negative for inconsistent ones (e.g. MT and color). See “Cortical organization analysis” section in SI Methods for details.

## Acknowledgements

We thank Andre Bastos, Mikael Lundqvist, Morteza Moazami, Jefferson Roy, Jason Sherfey, and Andreas Wutz for helpful discussions, and Mattia Rigotti for providing code and advice on the dimensionality analysis. This work was supported by grant NIMH 5R37MH087027 (E.K.M), grant ERC StG335880 (M.S.), and the Centre for Integrative Neuroscience (DFG, EXC 307) (M.S.).

## Supplementary Information

### SI Methods

#### Electrophysiological data collection

Parylene-coated tungsten electrodes were acutely inserted into target brain regions each day using a manual microdrive system. Neuronal activity was referenced to animal ground, amplified by a high-impedance headstage, filtered to extract spiking activity, digitized, and streamed to disk by an integrated electrophysiological system (Recorder or Omniplex, Plexon). The filtered signal was threshold-triggered to separate neuronal spikes from background noise, and individual spike waveforms were manually sorted offline into isolated neurons (Offline Sorter 3, Plexon). Neurons were included in analyses only for the duration of time in which they were well isolated from background noise and other neurons.

#### Behavioral paradigm

Two adult rhesus macaques *(Macaca mulatta),* one male and one female, were trained to perform a cued multidimensional categorization task. All stimuli were displayed on a color-calibrated CRT monitor at 100 Hz vertical refresh rate. An infrared-based eye-tracking system (Eyelink II, SR Research) continuously monitored eye position at 240 Hz. Behavioral control was handled by the MonkeyLogic program (www.monkeylogic.net) (35).

Each trial (Fig. 1A) was initiated when the animal fixated on a central dot (± 1.2° visual angle). Following a 500 ms fixation period, a centrally-presented visual cue (1000 ms) instructed the monkey to perform one of two task rules: color categorization (“greenish” vs. “reddish”) or motion categorization (“upward” vs. “downward”) of a subsequent centrally-presented colored, moving random-dot stimulus. Within each stimulus, all dots had the same color and moved in the same direction (100% coherence). The monkeys reported the stimulus category via a saccade towards a target to the left or right. The stimulus-response mapping for each task rule was fixed (color rule: greenish→left, reddish→right; motion rule: upward→left, downward→right). The monkeys were free to respond at any point (up to 3 s) after the random-dot stimulus onset.

The cues were four different gray shapes; two of them instructed the color rule, while the other two instructed the motion rule (Fig. 1B). Using two distinct cues for each rule allowed dissociation of neural activity related to the visual cue shapes and the task they instructed. Each motion category consisted of two directions (Fig. 1C; upward: 90°, 30°; downward: −30°, −90°), and each color category consisted of two hues (greenish: 90°, 30°; reddish: −30°, −90° hue angle; colors were defined in CIE L*a*b* space and had the same luminance and saturation). Additional tested values on one or more category boundaries were not included in the present analyses, as we focus here on unambiguously classifiable stimuli.

The monkeys performed both categorization tasks with high accuracy: 94% correct on average for motion, 89% for color. Across all trials and sessions in the current dataset, the median reaction time was 274 ms, with only 1% of all responses occurring earlier than 200 ms.

#### Data analysis

##### General

For most analyses, spike trains were converted into smoothed rates via convolution with a Hann function (half-width 125 ms; nearly equivalent to a 50 ms SD Gaussian, but with finite extent). To summarize results, we pooled rates or other derived statistics within empirically-defined epochs of interest for each task domain and area. The start of each epoch was set by the onset latency of total explained variance for a given task variable and area (see “Variance-partitioning model” section below), as estimated by the time point where the population total variance first attained 25% of its maximal value across all time points. To capture the full temporal extent of relevant neural activity, without introducing confounding factors, the ends of cue/task epochs for all areas were set at the onset of the random-dot stimulus, and the ends of motion- and color-related epochs were set at the earliest 1 percentile of the distribution of all behavioral reaction times (200 ms). All analyses were performed using custom Matlab code.

##### Hypothesis testing

All hypothesis tests used non-parametric bootstrap methods that do not rely on specific assumptions about the distributions of data values, unless otherwise noted. One-sample tests (Fig. 3B,E; 4B,E; 5B,E) resampled the statistic of interest with replacement 10,000 times from the neuronal population. The resulting distribution, offset by the actual observed statistic value, is an empirical estimate of the distribution of the statistic under the null hypothesis, and the one-tailed significance level was computed as the proportion of resampled values greater than the actual observed value. Two-sample tests (Fig. 3C,F; 4C,F; 5C,F) were computed similarly, but with separate bootstrap resampling from each compared population, evaluating the difference in the statistic of interest between populations, and computing the twotailed significance level as the proportion of absolute resampled values exceeding the absolute observed value. All presented standard errors were estimated as the 68% confidence intervals across bootstrap samples.

##### Variance-partitioning model

Our primary goal was to characterize the categoricality of the population of neurons in each cortical region—the degree to which they reflected the raw sensory stimuli or their abstracted categorical meaning. We quantified this by fitting each neuron’s spike rates, at each time point, with a linear model that partitioned rate variance across *n* trials into between-category and within-category effects:

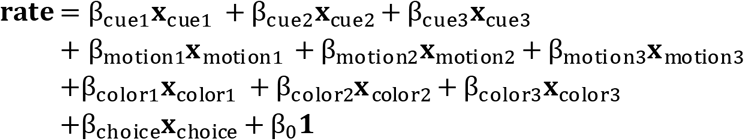

In this equation, 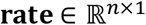 is a trial-length vector of spike rates for a given neuron and time point. For each task domain (rule cues, motions, and colors), the model included three orthogonal contrast terms (Fig. 2A). The first term **x**_*1_ ∈ {−1, 1}^*n*×1^ (where * indicates cue,motion, or color) contrasted the two stimulus items (cue shapes, motion directions, or colors) in one categorical grouping (rule, motion category, or color category) against the two items in the other categorical grouping ([A or B] – [C or D] in Fig. 2A). That is, it was a length *n* vector with a value of 1 for each trial where the stimulus item was A or B, and −1 for each trial where the stimulus was C or D. Thus, it reflected the actual task-relevant grouping of stimulus items into categories, and its associated fitted scalar coefficient 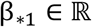 captured between-category variance. The other terms **x**_*2_, **x**_*3_ ∈ {−1, 1}^*n*×1^ were contrasts between the two other possible, non-task-relevant paired groupings of items ([A or C] – [B or D] and [A or D] – [B or C] in Fig. 2A). Thus, their associated coefficients 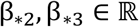 together captured within-category variance. The model also includes a term described below that accounted for behavioral choice (**x**_choice_), and a constant term **1** ∈ {1}^*n*×1^ (i.e. an all-ones vector) and associated coefficient (β_0_) to account for the overall condition-independent mean rate. The effect of each model term was quantified in terms of its percent of total data variance explained, measured from the difference in residual variance between the full model and a reduced model with the given term deleted (12). Note that this is an extension of the analysis strategy used to model task rule coding in our previous publication (11) to all three categorical domains in this task.

Across-trial mean neural responses for the four stimulus items in each domain (task cues, motions, or colors), for each neuron and time point, can be considered as four-dimensional vectors within the four-dimensional vector space 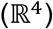 of all possible sets of item responses. If the three contrast terms described above are also considered as four-dimensional vectors (i.e. for items [A, B, C, D], **x**_*1_ would be [1 1 −1 −1], **x**_*2_ would be [1 −1 1 −1], and **x**_*3_ would be [1 −1 −1 1]), then along with the constant term ([1 1 1 1]), they constitute an orthogonal basis for this space. (This follows from the fact that they are mutually orthogonal and contain the same number of vectors as the dimension of the space (36)). This entails that any possible set of neural responses to the four items can be expressed as a linear combination of these terms. Hence, they capture all variance reflecting cue, motion, or color selectivity in the task. Since the first term (**x**_*1_) perfectly captures between-category variance, it follows that the other terms together capture all within-category variance. Thus, the three contrast terms partition the total variance in each task domain into between-category and within-category effects.

Within the context of each task rule, motion and color categories were—by design—inextricably linked with the monkey’s behavioral choice (e.g., for the color rule, greenish and reddish colors always mandated leftward and rightward saccades, respectively). For this reason, it is effectively impossible to dissociate category and choice effects within the context of each individual task rule, and to directly compare category effects across the two rules. When both rules are considered together, this identity link is broken, permitting dissociation of choice and category effects. However, there remains a partial (50%) correlation between choice and stimulus category. Therefore, it was critical to ensure that any activity reflecting choice (or subsequent motor preparation processes) was not spuriously interpreted as categorical coding. To partition out choice effects, we also included an additional model term **x**_choice_ ∈ {−1, 1}^*n*×1^ contrasting the two possible behavioral choices, with a value of 1 for rightward and −1 for leftward saccades. Its associated coefficient β_choice_ accounted for any variance due to choice, and thus category effects are measured in terms of their additional variance explained once choice effects are already accounted for (12). Choice effects were analyzed in detail in our previous publication on this dataset (11), and were therefore not examined further in this work.

Each model, 11 parameters in total, was fit via ordinary least squares separately for each neuron and time point. We used the bias-corrected ω^2^ formulation for explained variance (37).

##### Categoricality index

For each task variable (task cues, motions, or colors), the sum of explained variances for all model three terms sets an upper bound for the between-category variance alone—they can be equal only for a perfectly categorical population with zero within-category variance (Fig. 2B, top; simulated in Supplementary Fig. S1A, right). A purely sensory-driven population would instead have equal variance for all three terms. For example, a neuron with a preferred response for only a single stimulus item would have equal variance for all three contrasts, as each item appears once in every contrast (simulated in Fig. S1A, left). A population of neurons with preferred responses for arbitrary pairs of stimulus items—without regard to the task-relevant groupings—would also have equal expected variance for all three contrasts (simulated in Fig. S1A, center). Thus, the mean of all three contrasts provides an expected lower bound for the between-category variance in purely sensory populations (Fig. 2B, bottom). (Note that, for populations specifically encoding within-category differences, between-category variance can be less than the mean within-category variance.)

To measure where neural populations fall between these extremes, we computed a “categoricality index” equal to the area between the between-category and sensory (lower-bound) time series, expressed as a fraction of the full area between the total (upper-bound) and sensory (lower-bound) time series (Fig. 2C).

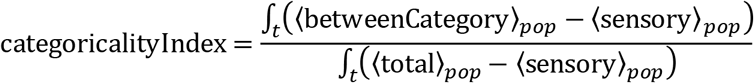

Here, betweenCategory = *ν*_1_ is the between-category (term 1) explained variance; total = *ν*_1_ + *ν*_2_ + *ν*_3_ is the sum of explained variances for all three terms; and sensory = (1/3)(*ν*_1_ + *ν*_2_ + *ν*_3_) is the mean of explained variances across all three terms. Each quantity is computed separately for each neuron and time point. The means 〈·〉_*pop*_ are over all sampled neurons in each area’s population, and the integrals ∫_*t*_(·) are over all time points within a pre-defined epoch for each area and task domain. This index is a specific measure of how categorical a neural population is, and ranges from 1 for a perfectly categorical population to 0 for a purely sensory population. (Negative values are possible if within-category variance is greater than between-category variance, i.e. for populations that specifically reflect within-category differences.)

In addition to the population categoricality index described above, we also computed indices for each individual neuron (Fig. S3). Here, all of the population means in the above equation were simply replaced by the corresponding quantities for each neuron. This analysis was restricted to neurons found to have significant total domain variance (*p* < 0.01; *F*-test), since the index is undefined—and the concept of categoricality is not meaningful—in the absence of any rate variance across the constituent items.

The statistic of category information used in our previous publication (11) was a debiased version of the between-category variance, equal to the between-category variance (*ν*_1_) minus the mean within-category variance ((1/2)(*ν*_2_ + *ν*_3_)). It can be shown that our categoricality index is equivalent to normalizing this statistic by the total domain variance (*ν*_1_ + *ν*_2_ + *ν*_3_):

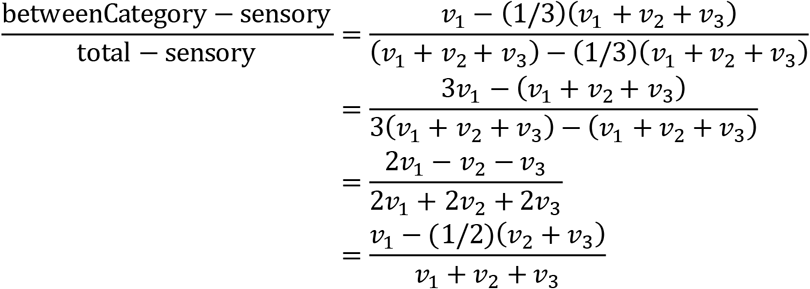

Thus, our index can also be viewed as a normalized contrast of between-category and within-category variance.

##### Neural simulations

To validate our analysis, we measured its properties on synthesized neural activity, where different coding attributes could be manipulated under experimental control. Note that the goal here is only to ensure the analysis behaves as expected in a simple, plausible toy model where ground truth is known, not to make any claims that our simulations accurately reflect actual cortical coding. We simulated populations of 200 neurons with Poisson rates modulated by both selectivity for categories and sensory tuning for stimulus items. Category selectivity in all task domains was simulated by a binary step function. A preferred Poisson rate was set for all items in one category, and a non-preferred rate for all items in the other category, with the preferred category randomized across simulated units. For task cues, stimulus tuning was simulated by setting a preferred rate for one or two cues randomly selected for each unit, without regard to categorical divisions. As a simple, plausible model of tuning for stimulus motion or color, we simulated each unit with a von Mises (circular Gaussian) tuning curve, with its center randomly oriented within the full range (0–360°) of motion directions or hue angles. In some simulations, we also modelled binary selectivity for behavioral choice (leftward vs. rightward saccades), with the preferred choice randomized across units.

Simulations evaluated four distinct measures of categorical coding: the normalized categoricality index introduced here (Fig. S1A–D), raw between-category variance (Fig. S1E,F), the debiased between-category variance statistic used in our previous publication (11) (Fig. S1G,H), and the category preference index (Fig. S1I,J).

In reported simulations, we expressed the tested selectivity in all population units, but manipulated its strength (the difference in Poisson rate between preferred and non-preferred items). Similar results were obtained if strength was fixed and the proportion of units expressing the tested selectivity was manipulated instead. All of the presented results also generalize well from the specific population size, tuning functions, proportion of units expressing selectivity, and response model assumed in these simulations. We evaluated simulated populations under the same task structure and analysis as the real data, and report means and standard deviations across 100 independent simulations.

These results are presented in the “Neural simulations” section of SI Results.

##### Category/choice consistency analysis

As an additional control to show our results are not confounded by behavioral choice, we compared color and motion category preferences under the two task rules, where category and choice coding make clear, opposing predictions. We fit models containing only the motion and color between-category terms, separately for the motion and color rules (variables here have the same interpretation as in equation 1):

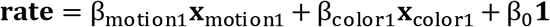

Two models were fit to each neuron’s spike rate pooled over the random-dot stimulus epoch. One model was fit to trials where the motion rule was instructed, another to trials where the color rule was instructed. For each term and task rule, we computed explained variance as above, but with a sign reflecting the preferred category: negative for neurons preferring downward motion or reddish color, and positive for upward motion or greenish color. Note that when a given task domain is task-relevant (i.e. motion during the motion rule, or color during the color rule), labeling by preferred category is equivalent to labeling by preferred choice (see “Variance-partitioning model” section above for details).

Choice coding predicts consistent motion and color category preferences (signs and magnitudes) when they are each task-relevant based on the currently instructed rule (i.e. when they both map to the same choice and resulting saccade direction). Category coding instead predicts consistent motion category preferences across both task rules, and the same for color category preferences. Consistency is visualized with scatterplots and quantified with Spearman rank correlation for each of these conditions. These results are presented in the “Category/choice consistency analysis” section of SI Results.

##### Preferred condition analysis

To qualitatively confirm our results using an alternative, more standard method of measuring category effects, we computed population average spike rates for preferred and non-preferred categories and stimulus items within each category. For each neuron and task variable, the preferred and non-preferred category (task rule, motion category, or color category) and the preferred and non-preferred constituent stimulus item within each category (rule cue, motion direction, or color) were determined from the average spike rate within the relevant time epoch for each area and task variable. Spike rate time series were averaged across the population of neurons in each area separately within each of the resulting four sorted conditions (preferred item in preferred category, non-preferred item in preferred category, preferred item in non-preferred category, non-preferred item in non-preferred category). To avoid circularity in this analysis, we used a condition-balanced ten-fold crossvalidation method. Preferred conditions were estimated from 90% of trials and the resulting sorting was used for population averaging of rate series across the remaining 10% of trials, with each partition having approximately balanced trial numbers across the four conditions. This was performed for each disjoint subset of 10% of trials, and the final presented results are the average of these ten cross-validation folds.

In this analysis, the signature of categorical coding is a large difference in rate between the preferred and non-preferred category, with minimal rate differences between items within each category. For sensory representations, rate differences between and within categories would be of similar magnitude. Therefore, to summarize these results we computed a category preference index, a contrast index comparing differences in preference-sorted mean rates between categories vs. within categories.

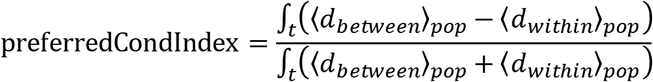

In this equation, *d_between_* is the difference in rate between the preferred category and the nonpreferred category for each neuron,

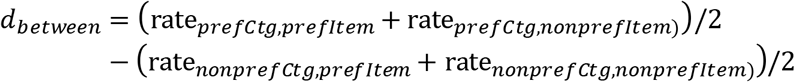

and *d_between_* is the difference in rate between the preferred item and the non-preferred item within each category, averaged across both categories, for each neuron.

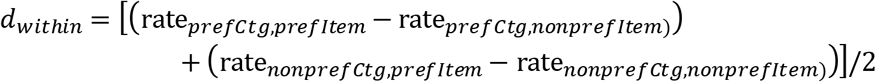

Both the between- and within-category differences were rectified at zero, as negative rate differences have little meaning in a preference-sorted analysis (these were rare, but possible because of the cross-validation procedure). These results are presented in the “Preferred condition analysis” section of SI Results.

##### Population dimensionality analysis

To measure the dimensionality of population activity, we adapted a method developed by Machens and colleagues (17), and extended by Rigotti and colleagues (18). This method estimates the dimensionality of the space spanned by population activity vectors by determining the number of principal components required to describe them. Spike rates were computed within each of the time epochs enumerated above, and the resulting data for each area was pooled across all electrodes and recording sessions into “pseudopopulations”. To extrapolate results to larger neural populations, we generated artificial neurons via specific relabeling of conditions, under the assumption that the full underlying population likely includes neurons with similar activity distributions, but slight differences in tuning for rule cues, motion directions, and colors (18). Labels were swapped between the two visual cues associated with each task, between the two colors within each color category, and/or between the two directions within each motion category. Note that these label reassignments both maintain the semantic logic of the paradigm and would not alter the information carried by either sensory or categorical neurons, as estimated above. Artificial neural populations were generated by all permutations of performing each of these relabeling operations or not, resulting in a 64-fold multiplication of the actual populations (2^2^ colors × 2^2^ directions × 2^2^ cues). Extrapolated populations of different sizes were generated by randomly subsampling from this full population of real and artificial neurons, separately for each cortical area. Dimensionality values obtained from analyzing only the actual recorded populations (Fig. 6A,C, squares) generally fell within the distribution of those obtained from the extrapolated populations, suggesting that this procedure did not substantially alter the results.

For this analysis, we characterized neural activity within the full space of 64 task conditions (4 rule cues × 4 motion directions × 4 colors) and within the random-dot stimulus epoch, when all 64 conditions were differentiated. An additional analysis was performed within the reduced space of 16 motion directions × colors. To avoid PCA being dominated by a few highly active neurons, trial spike rates for each neuron were preprocessed by *z*-scoring them by the mean and SD across all trials, with a regularization constant of 0.1 added to each SD (18). For each area and extrapolated population size, the vector of mean spike rates across all neurons for each condition was computed, and the dimensionality of the space spanned by each resulting set of 64 neural population activity vectors was estimated as the number of their principal components (eigenvalues) significantly greater than the estimated distribution of principal components due to finite sampling noise. Noise was estimated by computing the difference between randomly chosen trials within each condition, weighted by its expected contribution to the estimated condition mean (17), and submitting the resulting set of 64 noise vectors to PCA. This procedure was repeated 1000 times, randomly sampling within-condition trial pairs each iteration, to generate a distribution of noise-derived eigenvalues. Dimensionality was estimated as the number of data eigenvalues exceeding 95% of the distribution of first (largest) eigenvalues computed from the sampled noise (18). This entire procedure was repeated 50 times, selecting with replacement a different random subsample of area neurons in each iteration, to estimate bootstrap standard errors of the resulting dimensionality statistic. Qualitatively similar results were obtained using the method of Rigotti et al. 2013 based on the number of binary classifications that can be successfully decoded from population activity vectors.

##### Cortical organization analysis

To quantitatively relate our results to classical models of large-scale cortical organization, we separately fit the 18 population summary values (6 areas × 3 task variables) of total domain explained variance (i.e., sensory information; Fig. 7A) and categoricality index (Fig. 7B) with a simple two-predictor linear model:

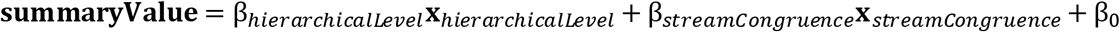

Here, **summaryValue** 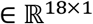 is the vector of 18 summary values—total domain variances or categoricality indices. The first predictor 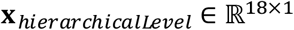 was the Felleman and Van Essen hierarchical level of each area (MT,V4: 5; PIT,LIP: 7; FEF: 8; PFC: 10), estimated from the relative distribution of feedforward-type vs. feedback-type anatomical connections between areas (29). Its associated coefficient β_*hierarchicalLevel*_ accounts for any effects of each area’s hierarchical level on the given summary value. The second predictor **x**_*streamcongruence*_ ∈ {—1,0,1}^18×1^ reflected the expected functional congruence of each combination of task variable and area based on the classical divisions of visual areas and functions of the ventral and dorsal processing streams. It took a value of 1 for consistent combinations (e.g. MT and motion, V4 and color), and −1 for inconsistent ones (e.g. MT and color, V4 and motion). For this purpose, rule cue (shape) and color were designated as ventral stream domains, and V4 and PIT ventral stream areas. Motion direction was designated a dorsal stream domain, and MT, LIP, and FEF dorsal stream areas. Since PFC is thought to integrate across both streams (29, 38), all variable/area combinations involving it were set to zero for this predictor. To equate the range of stimulus information and categoricality index values across task variables, values across allareas for each variable were normalized to range from 0 to 1. Prior to this normalization, categorical information values were rectified at zero, thus equating areas that match the sensory-based prediction with those less than it (reflecting slight biases toward within-category selectivity). Each model (3 parameters in total) was fit via ordinary least squares, and evaluated by the variance explained by each of the two predictors. Errors and significance levels were estimated by bootstrapping these statistics over the contributing neurons 10,000 times.

### SI Results

#### Neural simulations

To confirm that our analysis methods performed as expected, we first assayed their properties on synthesized neural activity with known ground truth (see “Neural simulations” section in SI Methods for details). The basic properties of all analyses considered here are summarized in Supplementary Table S1. We first confirmed the categoricality index was ≈ 1 for populations of units with ideal binary category selectivity for task rules (Supplementary Fig. S1A, right), and was ≈ 0 for populations with only sensory selectivity for single task cues (Fig. S1A, left) or for random pairs of cues (Fig. S1A, center). To examine this in more detail, we parametrically varied both the relative strength of simulated sensory and categorical signals (Fig. S1B–C, *x*-axis), and the overall strength of both signals in concert (i.e. simulated differences in spike rates between preferred and non-preferred conditions; Fig. S1B–C, color saturation). For task rule coding, we simulated sensory selectivity for random pairs of cues (Fig. S1B; results are similar for single-cue selectivity). For motion coding, we simulated Gaussian sensory tuning for motion direction (Fig. S1C; note that color and motion are treated identically in these simulations and results therefore generalize to both domains). These simulations confirmed that our categoricality index reliably reflected sensory/category relative strength (curves monotonically increase from left to right), but was relatively invariant to overall signal strength (different-color curves largely overlap). Finally, to confirm that our modeling procedure successfully partitioned category and choice effects, we simulated populations independently varying in both the relative weight of sensory and categorical motion signals (Fig. S1D, *x*-axis), and the strength of choice signals (Fig. S1D, color saturation). These simulations confirmed the categoricality index was also relatively insensitive to choice signal strength. As a positive control, we demonstrated that the variance explained by choice itself did indeed robustly reflect choice signal strength (Fig. S1D, inset).

Raw between-category variance, in contrast, provides a biased and inconsistent measure of categoricality. As expected, between-category variance was high for a simulated population with ideal binary category selectivity for task rules (Fig. S1E, right). However, it also had non-zero values for populations with sensory selectivity for single task cues (Fig. S1E, left) or for random pairs of cues (Fig. S1E, center). Figure S1F shows that between-category variance conflates sensory/categorical relative strength with overall spike rates. Even purely sensory populations (Fig. S1F, leftmost points) have non-zero values that increase proportionately to total signal strength. Results are similar for simulated sensory selectivity for single task cues, and for motion direction and color. Thus, the raw percent of variance explained by categories is an inadequate measure of population categoricality.

In a previous publication, we proposed a debiased between-category variance statistic of categorical task rule information (11). This measure subtracts within-category variance from the raw between-category variance (see “Categoricality index” section in SI Methods for details), resulting in an expected value of zero for purely sensory representations. This was confirmed using the same set of simulations as above (gray bars in Fig. S1G; leftmost points in Fig. S1H). However, for populations with a mixture of sensory and categorical coding (right portion of Fig. S1H), this statistic is influenced by the overall signal strength. For any given value of sensory/categorical relative weight (*x*-axis value) it is proportional to overall rate, and any given value of this statistic (*y*-axis value) might correspond to a wide range of combinations of categoricality and spike rate. Similar results obtained for other sensory domains. Thus, while this statistic provides a measure of category information that is unbiased for purely sensory activity, it does not in general unambiguously report population categoricality, as defined here.

As an alternative method to measure population categoricality, we computed population average spike rates for preferred and non-preferred categories and within-category stimulus items. We summarized this analysis with a category preference index, a contrast index contrasting between-category and within-category differences in the preference-sorted mean spike rates (see “Preferred condition analysis” section in SI Methods for details). Like the categoricality index, this statistic reports values ≤ 0 for sensory populations (Fig. S1I, left and center) and ~ 1 for categorical populations (Fig. S1I, right), and is relatively invariant to overall signal strength (Fig. S1J).

#### Category/choice consistency analysis

As an additional control to show our results are not confounded by behavioral choice, we compared color and motion category preferences under the two task rules, where category and choice coding make clear, opposing predictions. Category preference was quantified by the between-category motor or color variance, with a sign reflecting the preferred category: negative for downward or reddish preferring, positive for upward or greenish preferring. Consider the comparison between motion category preference during the motion rule vs. color category preference during the color rule, i.e. when each category is task-relevant and drives behavioral choice (Fig. S5A and S5D–E, left). A neuron encoding choice would appear to have the same sign and magnitude of category preference for both conditions, as these would map to the same choice (saccade direction). A population dominated by choice selectivity would thus be expected to have a strong consistency of signed variance between these conditions, and we therefore refer to this comparison as “choice-consistent”. In contrast, category-selective populations would be expected to either carry information about only a single domain or have unrelated preferences across category domains, and thus have little consistency between these conditions.

Now consider instead the comparison between motion category preference during the motion rule vs. during the color rule (Fig. S5B; Fig. S5D–E, center), or between color category preference during the motion rule vs. during the color rule (Fig. S5C; Fig. S5D–E, right). A neuron encoding a given category across both rules would have the same sign and magnitude of preference in both conditions, and a population dominated by category selectivity would thus show strong consistency between these conditions. We therefore refer to these comparisons as”category-consistent”. In contrast, a choice-selective neuron would appear to carry category information only when the category was task-relevant, and a choice-dominated population would thus have inconsistent category preferences across task rules.

To determine the type of coding that is dominant overall in each area, we plotted scatterplots and computed Spearman rank correlations across the entire sampled neural populations for each comparison above. For the choice-consistent comparisons, the scatterplots showed no obvious relationship between motion and color category preference (Fig. S5A), and their correlations were weak (Fig. S5D, left) and only significant for V4 (*p* = 0.002; *p* ≥ 0.01 for all other areas, permutation test). In contrast, the category-consistent comparisons had clear structure (Fig. S5B,C) and stronger correlations (Fig. S5D, center and right), which were significant for at least one category domain in all areas (*p* < 0.01, Bonferroni-corrected for two comparisons). This suggests that category signals are generally dominant over choice signals in the studied areas. Note that PFC, FEF, and to some extend LIP did have choice-consistent and category-consistent correlations of similar magnitude. This suggests some heterogeneous mixture of category and choice effects in these populations, consistent with previous reports showing strong choice and saccade signals in these areas (11, 39, 40).

As a final check that our main analysis specifically extracts category effects from these population mixtures, we repeated the correlation analysis for only the subset of neurons carrying significant category information across the full set of trials (*p* < 0.01, F-test on between-category variance for equation 1 model fit to stimulus epoch rates across both rules; Fig. S5A-C, colored dots). There is some circularity in this analysis, in that the criterion used to select neurons (significant category variance across all trials) is not entirely independent of the measure examined (signed category variance under each rule). However, we felt it was helpful to include these results to provide a complete picture of the data and analysis. For the subset of category-selective neurons, choice-consistent correlations remained weak (Fig. S5E, left) and were not significant any area (*p* ≥ 0.01), while correlations for the category-consistent conditions were uniformly strong (Fig. S5E, center and right). Together, these results provide another line of evidence that our main analysis correctly measured category signals, rather than confounded choice signals.

#### Preferred condition analysis

To confirm our results using an alternative method of measuring category effects, we computed population average spike rates for preferred and non-preferred categories and stimulus items within each category (Fig. S6). These results were summarized with a category preference index, a contrast index comparing between-category to within-category differences in mean rates (see “Preferred condition analysis” section in SI Methods for details). The results indicated broad agreement with the categoricality index in the main text. For task cue coding in PFC and FEF, rate differences between the preferred and non-preferred task rules were considerably larger than between cues instructing the same rule (Fig. S6A), and their category preference indices were significantly greater than zero (Fig. S6B). LIP trended in the same direction, but unlike the main text analysis (Fig. 3E in the main text), it was not significant. For MT, V4, and PIT, between- and within-category differences were similar (Fig. S6A), and their indices were not significantly different from zero (Fig. S6B). For motion coding, only PFC, FEF, and LIP showed significant category preference indices (Fig. S6E). Again, this result was overall similar to the main text categoricality results, though in that case FEF failed to reach significance (Fig. 4E in main text). As for the categoricality index analysis (main text Fig. 5E), color coding exhibited a more categorical representation overall, with all areas except MT having significant category preference indices (Fig. S6H). Note that perfect alignment between results from these two methods would not be expected, due to their many salient differences (Supplementary Table S1). In the preferred condition analysis, choice effects were not partitioned out, rate was not normalized (and thus high-rate neurons might contribute disproportionately), and overall domain variance was not normalized out. The fact that results were, nevertheless, generally quite similar with an arguably more standard method adds support to our main conclusions.

#### Comparison with our previous results

Some of our results appear to be somewhat at odds with our prior publication on this dataset (11). In particular, it was previously claimed that V4 and PIT contained strong task rule signals (Fig. 2C in (11)), whereas we claim no significant task rule categoricality in these same areas (Fig. 3D,E). In part, this is due to differences in the studied neural signals—we analyzed isolated single neurons, whereas the previous study analyzed multi-unit signals. We repeated our analysis on multi-unit signals similar to those used previously (pooling together all threshold-crossing spikes on each electrode). Though results were generally quite similar (Fig. S4), multiunits produced a slightly more categorical rule cue representation in V4 and PIT (Fig. S4A), perhaps suggesting stronger signals for task rules exist in the smaller neurons likely to appear only in multi-unit signals.

The primary difference, though, is in the specific questions addressed by each study and their resulting analytical strategies. Our previous study addressed the overall task rule information conveyed by each neural population. Thus, it used a debiased category variance statistic, equal to the difference between between-category variance and the mean within-category variance. This statistic is unbiased for purely sensory representations—it has an expected value of zero (simulated in Fig. S1G). But for populations with any mixture of category effects, it reflects both categoricality and total domain variance (simulated in Fig. S1H). Thus, it can be high for populations—like rule cue representations in V4 and PIT—with strong total domain variance, even if only a very small fraction of that variance is categorical. As expected, recomputing our main findings with this statistic produced quite different results from ours (Supplementary Fig. S7), which qualitatively appear to look like a mixture of categoricality (Fig. 3E,4E,5E) and total domain variance (Fig. 3B,4B,5B). Here, we instead addressed how *categorical* neural representations are, independent of their overall information. Our categoricality index measures this (Supplementary Fig. S1B,C) by normalizing total domain variance out of the above statistic. Under this statistic, V4 and PIT contain task cue representations that are only weakly categorical (Fig. 3D,E). Thus, we can reconcile results from the two studies by concluding that V4 and PIT contain strong information about task cues but only a small fraction of that information is categorical. In contrast, despite the weaker overall task cue information in PFC and FEF, a substantial fraction of that information reflects the learned task rule categories. This definition accords well with both intuitive notions of categoricality and those previously proposed (3, 10).

### Supplementary tables

**Table S1.**
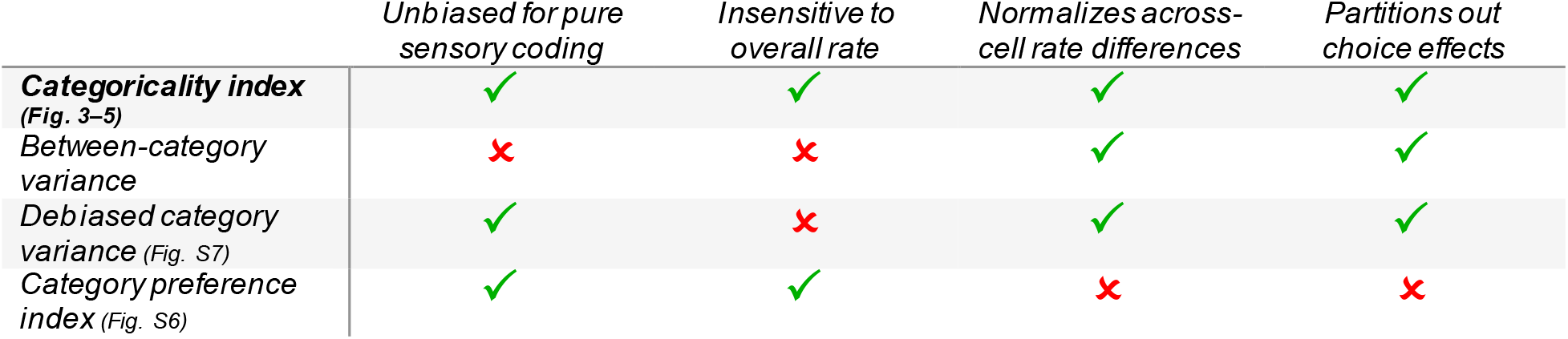
Comparison of properties of analysis methods used. Table compares basic properties of all analysis methods examined in this paper. A green check indicates the given analysis features the given property; a red x indicates it does not. Analyses listed in table rows are: the categoricality index, proposed in this paper (main text Fig. 3E,4E,5E; simulations in Fig. S1A-D); raw between-category variance (main text Fig. 3D,4D,5D; simulations in Fig. S1E,F); the debiased category variance statistic, proposed in Siegel et. al 2015 (Fig. S7; simulations in Fig. S1G,H); the category preference index (Fig. S6; simulations in Fig. S1I,J).

|  | Unbiased for pure sensory coding | Insensitive to overall rate | Normalizes across-cell rate differences | Partitions out choice effects |
| --- | --- | --- | --- | --- |
| **Categoricity index (Fig. 3-5)** | ✓ | ✓ | ✓ | ✓ |
| **Between-category variance** | ✗ | ✗ | ✓ | ✓ |
| **Debiased category variance (Fig. S7)** | ✓ | ✗ | ✓ | ✓ |
| **Category preference index (Fig. S6)** | ✓ | ✓ | ✗ | ✗ |

### Supplementary figure legends

**Figure S1.**
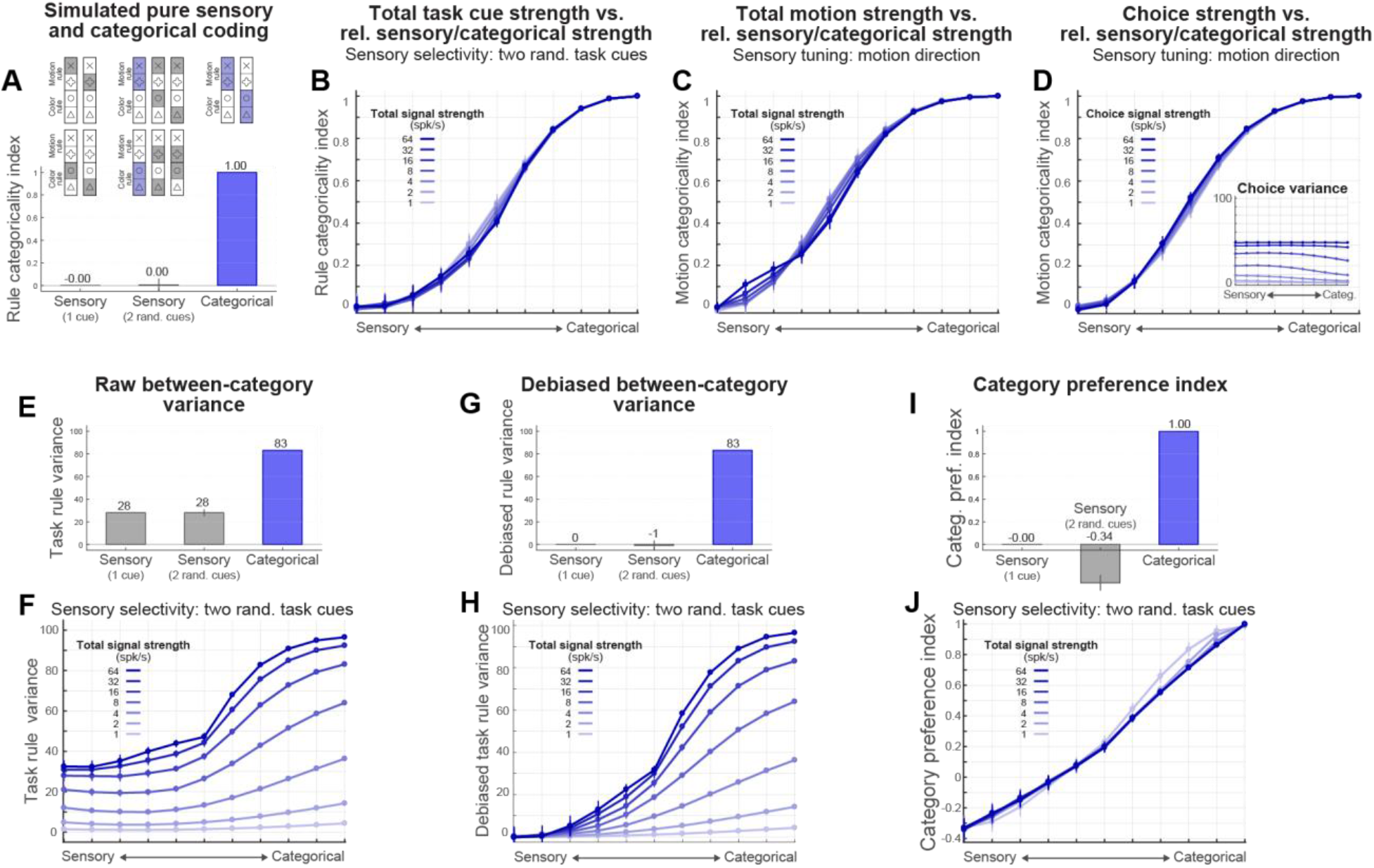
Validation of analysis methods with neural simulations. (*A*) Mean (± SD across independent simulations) categoricality index computed on simulated neural populations. Diagrams at top reflect simulated patterns of neural activity across the four task cues—white indicates non-preferred responses, blue indicates preferred responses consistent with task rule coding, and gray indicates preferred responses inconsistent with task rules. Results for three simulated populations are shown: one with sensory selectivity for single task cues (left), one with sensory selectivity for random pairs of cues (center), and one with pure categorical selectivity for pairs of cues that map to the same task rule (right). The categoricality index veridically reports values of ~0 for the two sensory populations and ~1 for the categorical population. (*B*–*C*) Mean (± SD) categoricality index for simulated populations parametrically varying in the relative strength of sensory and categorical signals (*x*-axis) and in the overall strength of both signals in concert (rate difference between preferred and non-preferred conditions; color saturation of plots). Results are shown for simulations with sensory selectivity for random pairs of task cues (*B*), and sensory tuning for motion direction (*C*). In all cases, index values reliably reflect sensory/categorical relative strength (curves monotonically increase from left to right), but are relatively insensitive to overall signal strength (different-color curves are mostly overlapping). (*D*) Mean (± SD) categoricality index for simulated populations parametrically varying in the relative strength of sensory and categorical motion signals (*x*-axis) and in the strength of signals for behavioral choice (rate difference between left and right saccades; color saturation of plots). Index values are not confounded by choice effects. Inset: The variance attributed to choice does reliably track choice signals, as expected. (*E*–*F*) Mean (± SD) raw task-rule between-category variance, under the same simulations as in *A* and *B*. Raw between-category variance has non-zero values for purely sensory populations (*E*, left and center) and is proportional to total signal strength (vertical shift of curves in *F*), indicating it is a biased measure of categoricality, which is confounded with global changes in spike rates. (*G*–*H*) Mean (± SD) debiased task-rule between-category variance, under the same simulations as in *A* and *B*. This is the statistic used to measure task rule information in our previous publication from this dataset (11), and is equal to between-category variance minus the mean within-category variance. This statistic is ~0 for purely sensory populations (*G*, left and center), indicating that it successfully removed the bias in the raw variance. However, for populations with a mixture of category effects, it reflects both categoricality and overall spike rates (vertical offset of curves in *H*). (*I*–*J*) Mean (± SD) task-rule category preference index, under the same simulations as in *A* and *B*. Like the categoricality index, this statistic reports values ≤ 0 for sensory populations (*I*, left and center) and ~ 1 for categorical populations (*I*, right), and is relatively invariant to overall signal strength (*J*).

**Figure S2.**
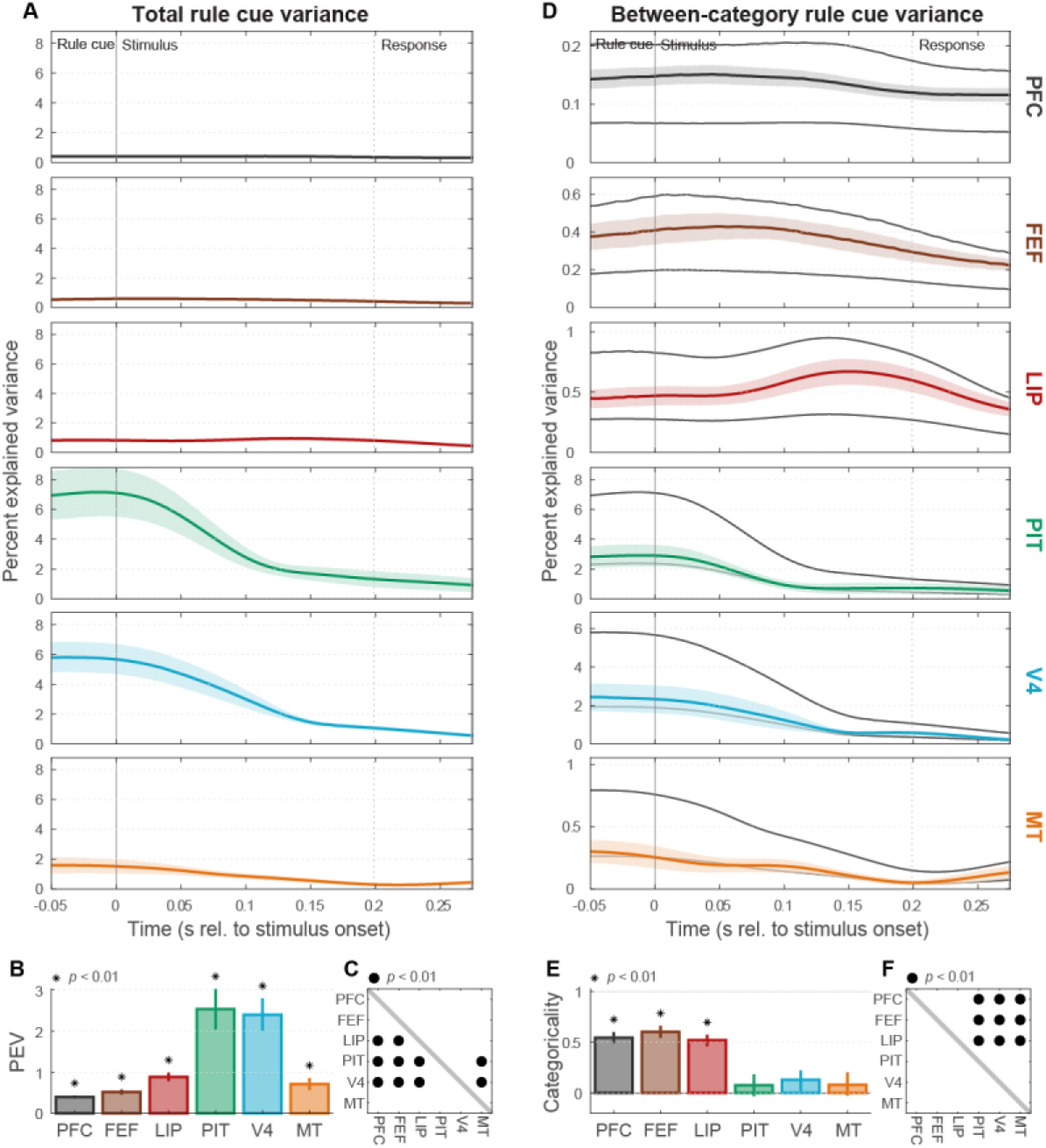
Task rule cue coding during stimulus period. (*A*) Population mean (± SEM) total rule cue variance (cf. Fig. 3A) during the random-dot stimulus period. Cue information in visual areas V4, PIT, and MT drops toward baseline soon after cue offset. (*B*) Summary (across-time mean ± SEM) of total stimulus-period rule cue variance for each area (cf. Fig. 3B). All areas retain significant cue information during the stimulus period (*p* < 0.01). (*C*) Indicates which regions (rows) had significantly greater cue information than others during the stimulus period (cf. Fig. 3C). (*D*) Mean (± SEM) between-category rule cue variance (task rule information). (*E*) Stimulus-period task rule categoricality index (± SEM) for each area (cf. Fig. 3E). PFC, FEF, and LIP remain significantly categorical (*p* < 0.01)—they continue to convey task rule information through the stimulus period. (*F*) Indicates which regions (rows) had significantly greater task rule categoricality indices than others (columns; *p* < 0.01; cf. Fig. 3F).

**Figure S3.**
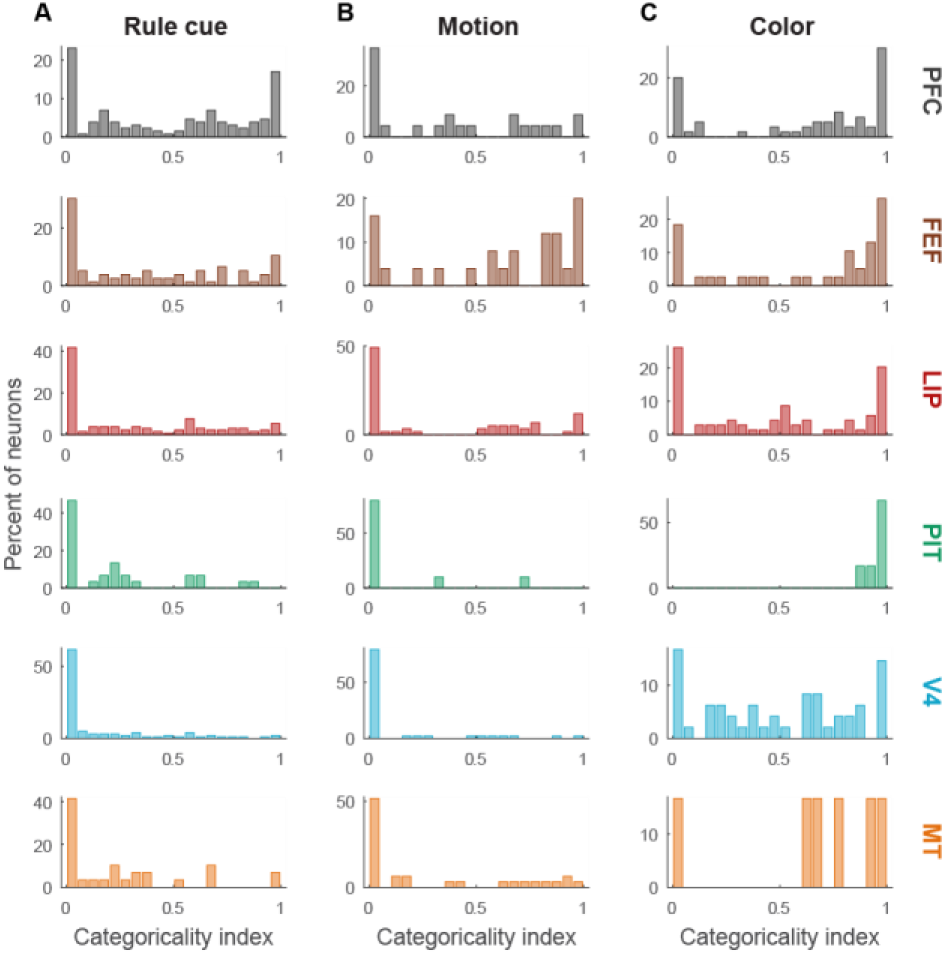
Distribution of single-neuron categoricality. (*A*) Population distributions of rule cue categoricality indices computed on single neurons in each area. (*B*) Population distributions of motion categoricality indices computed on single neurons in each area. (*C*) Population distributions of color cue categoricality indices computed on single neurons in each area. For all three domains, areas with high population categoricality indices (cf. Fig. 3E,4E,5E) have a large proportion of nearly purely categorical single neurons (index ~ 1). However, in almost all cases (except for PIT color coding) there also remains a residual subpopulation of sensory-driven neurons (index ~ 0), as well as single neurons whose activity reflects a mixture of between-category and within-category effects (0 < index > 1).

**Figure S4.**
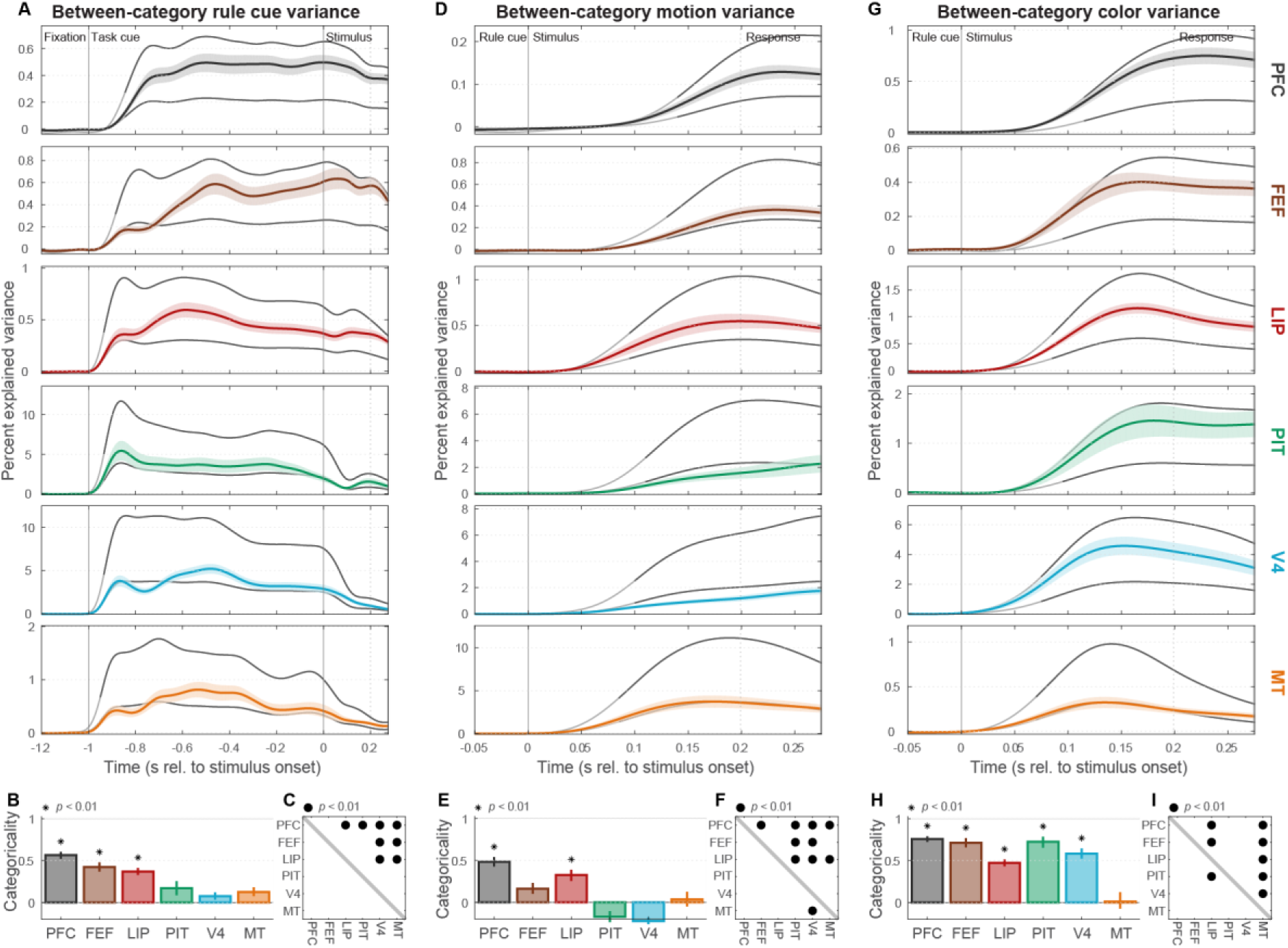
Main results are similar for multi-unit signals. (*A*) Population mean (± SEM) multi-unit between-category rule cue variance (cf. Fig. 3D). Multiunit signals were computed by pooling together all threshold-crossing spikes on each electrode. (*B*) Summary (± SEM) of multi-unit rule cue categoricality index (cf. Fig. 3E). (*C*) Cross-area significance matrix for multi-unit rule cue categoricality indices (cf. Fig. 3F). (*D*) Population mean (± SEM) multi-unit between-category motion variance (cf. Fig. 4D). (*E*) Summary (± SEM) of multi-unit motion categoricality index (cf. Fig. 4E). (*F*) Cross-area significance matrix for multi-unit motion categoricality indices (cf. Fig. 4F). (*G*) Population mean (± SEM) multi-unit between-category color variance (cf. Fig. 5D). (*H*) Summary (± SEM) of multi-unit color categoricality index (cf. Fig. 5E). (*I*) Cross-area significance matrix for multi-unit color categoricality indices (cf. Fig. 5F).

**Figure S5.**
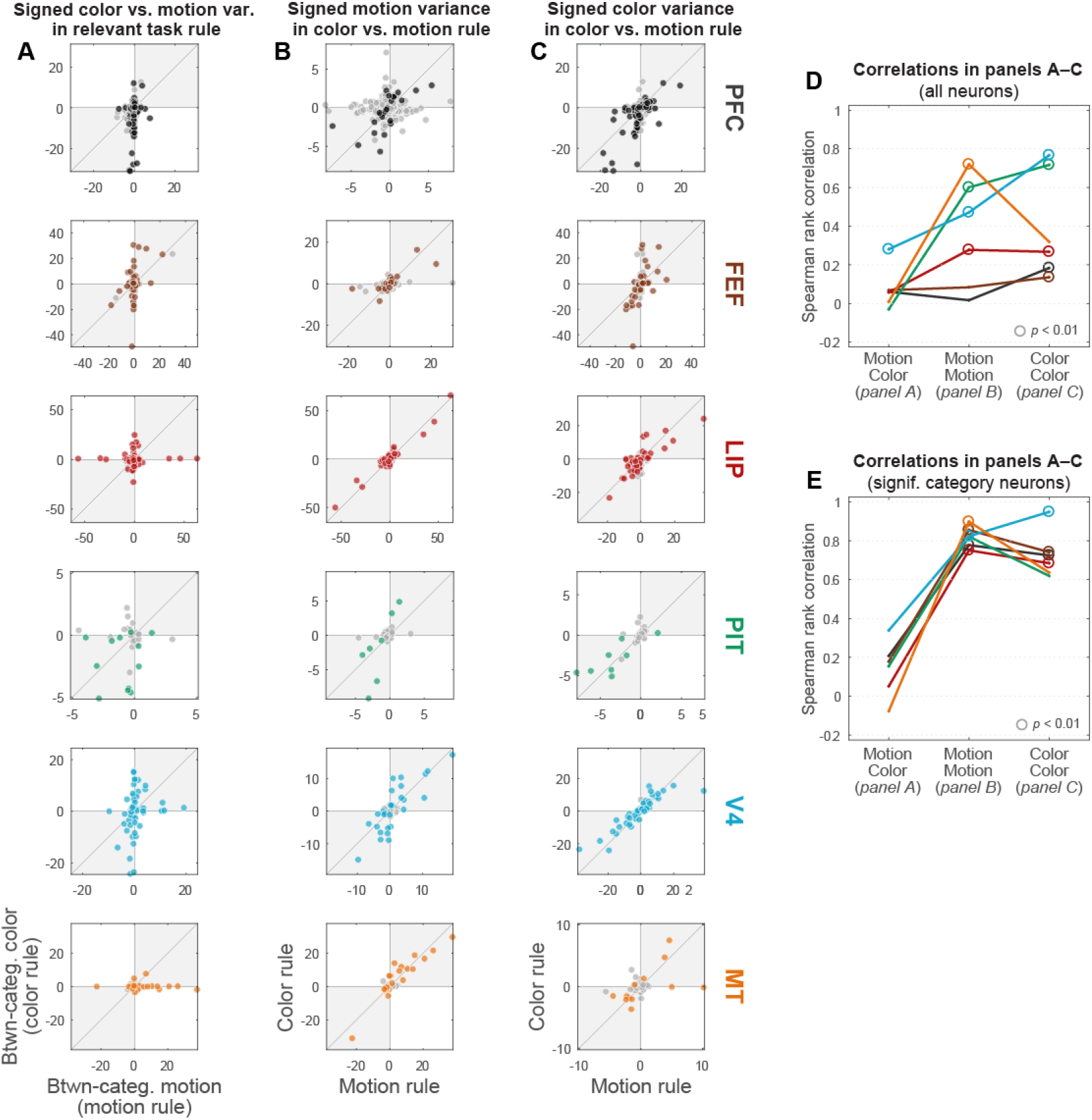
Category/choice consistency analysis. (*A*) Signed motion variance for each area neuron under the motion rule (*x*-axis) vs. color variance under the color rule (*y*-axis), i.e. when each category domain is task-relevant and drives behavioral choice. The sign for each neuron’s data point reflects its preferred category— negative for downward-preferring or reddish-preferring, and positive for upward-preferring or greenish-preferring. Neurons with significant between-category motion or color variance (*p* < 0.01; F-test) from the full-session analysis are colored in, while non-significant neurons are light gray. Neurons coding for behavioral choice (or subsequent motor preparation processes) would appear to have consistent motion and color category preferences, and thus would lie near the positive diagonal. (*B*) Signed between-category motion variance, measured separately within the motion (*x*-axis) and color rule (*y*-axis) trials. Signs reflect preferred motion categories—negative for downward-preferring, and positive for upward-preferring. Neurons with significant motion variance are colored in. Neurons coding for the same motion category irrespective of the task rule in effect would have consistent values and lie near the positive diagonal. (*C*) Signed between-category color variance, under the motion (*x*-axis) and color (*y*-axis) rules. Signs reflect preferred color categories—negative for reddish-preferring, and positive for greenish-preferring. Neurons with significant color variance are colored in. Neurons coding for the same color category irrespective of task rule would have consistent values and lie near the positive diagonal. (*D*) Spearman rank correlations for all neurons in each scatterplot in panels *A*–*C* (*x*-axis). Circles indicate significant correlations (*p* < 0.01; permutation test). Choice coding predicts stronger correlations for the motion/color (left, panel *A*) condition. Categorical coding predicts stronger correlations for the motion/motion (center, panel *B*) and color/color (right, panel *C*) conditions. Results are consistent with categorical coding dominating the overall population in most studied areas, except for PFC and FEF, which appear to contain a heterogeneous mixture of category and choice effects. (*E*) Correlations for only significant categorical neurons in each scatterplot in panels A-C (x-axis). Predictions are same as in D. All areas exhibit correlation patterns consistent with categorical, rather than choice, coding. This confirms that our variance-partitioning model successfully recovers categorical coding, unconfounded by choice signals.

**Figure S6.**
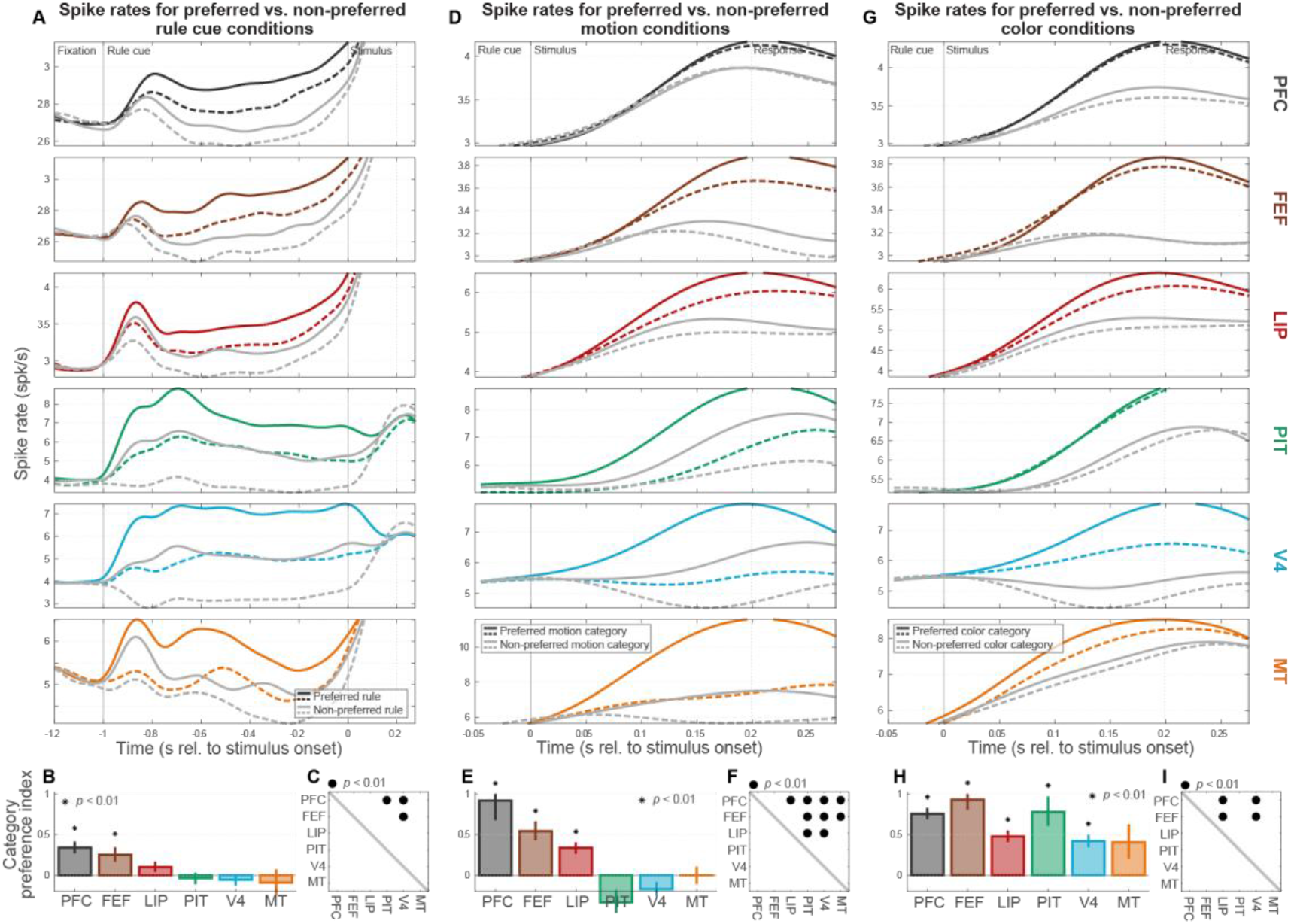
Preferred condition spike density analysis. (*A*) Population mean spike rates for preferred (colored lines) and non-preferred (light gray lines) task rules, and for preferred (solid lines) and non-preferred (dashed lines) rule cues within each task rule. This is plotted separately for each studied area as a function of within-trial time (referenced to the onset of the random-dot stimulus). Distinct subsets of trials were used to estimate preferred conditions and compute mean rates with the estimated sorting, to avoid circularity in the analysis. Note that rates for some areas exceed the plotting range outside the time epoch of interest here (the rule cue period). Broadly consistent with the analyses in the main text, rate differences for cues instructing the same task were small compared to differences between tasks in PFC and FEF, whereas for visual areas MT, V4, and PIT these differences were of comparable size. (*B*) Task rule category preference index (± SEM) for each area, which summarizes results in (*A*) by contrasting between-category and within-category population rate differences (cf. Fig 3E in the main text). Only PFC and FEF had index values significantly greater than zero (*p* < 0.01, asterisks). (*C*) Indicates which regions (rows) had significantly greater task rule preferred condition indices than others (columns; *p* < 0.01, dots; cf. main text Fig. 3F). (*D*) Mean spike rates for preferred (colored lines) and non-preferred motion categories (light gray lines), and for preferred (solid) and non-preferred (dashed) directions within each motion category. Between-category rate differences were large compared to within-category differences in PFC, FEF, and LIP, whereas for visual areas MT, V4, and PIT these differences were comparable. (*E*) Motion category preference index (± SEM) for each area, summarizing results in (*D*). (cf. main text Fig 4E). PFC, FEF, and LIP had index values significantly greater than zero (*p* < 0.01, asterisks). (*F*) Indicates which regions (rows) had significantly greater motion preferred condition indices than others (columns; *p* < 0.01, dots; cf. main text Fig. 4F). (*G*) Mean spike rates for preferred (colored lines) and non-preferred color categories (light gray lines), and for preferred (solid) and non-preferred (dashed) colors within each color category. Between-category rate differences were large compared to within-category differences in FEF, PIT, and PFC, but less so for areas MT, V4, and LIP. (*H*) Color category preference index (± SEM) for each area, summarizing results in (*G*). (cf. main text Fig 5E). FEF, PIT, PFC, LIP, and V4 all had index values significantly greater than zero (*p* < 0.01, asterisks). (*I*) Indicates which regions (rows) had significantly greater color preferred condition indices than others (columns; *p* < 0.01, dots; cf. main text Fig. 5F).

**Figure S7.**
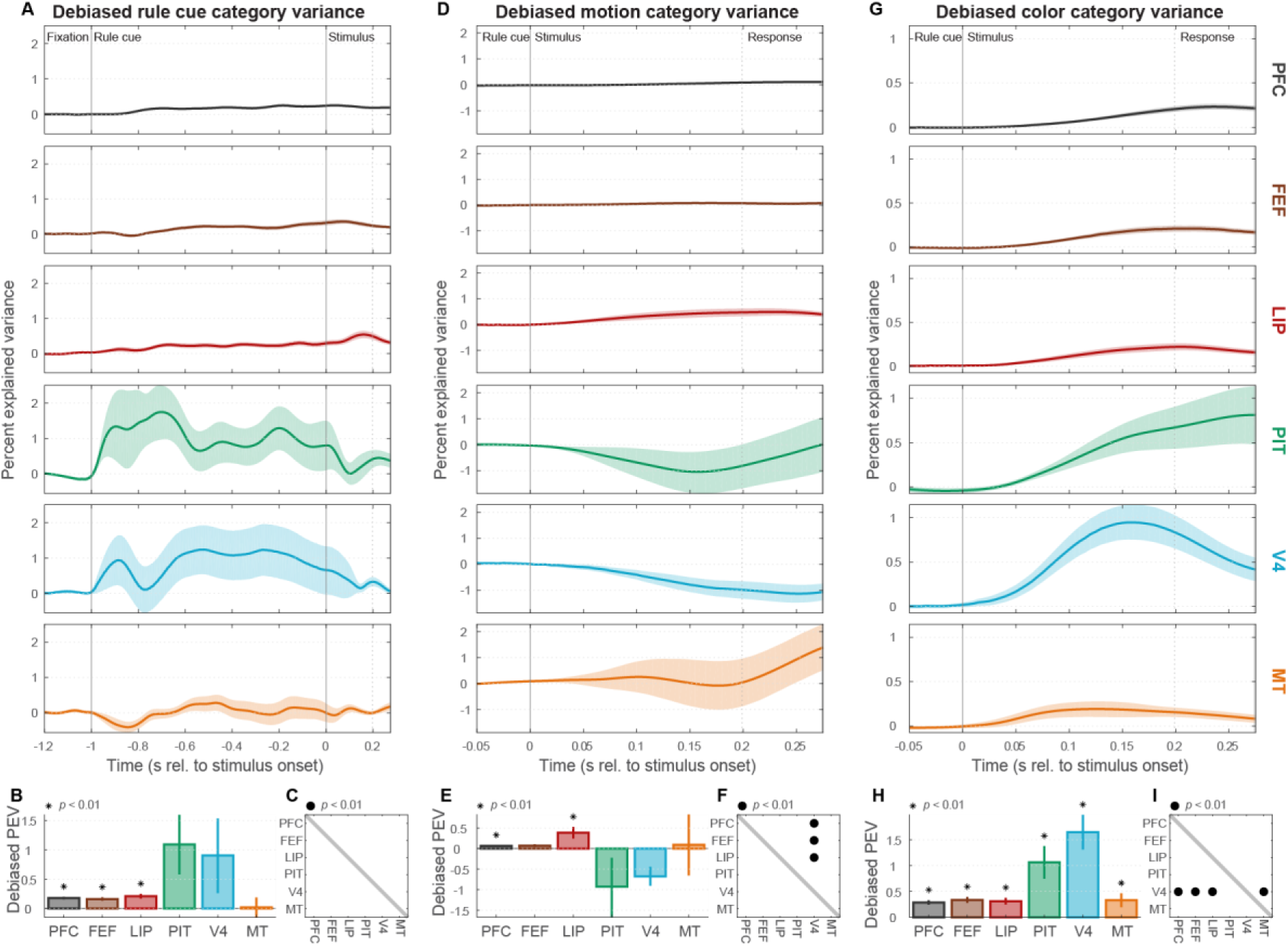
Debiased category variance results (cf. Siegel et al. 2015) (*A*) Population mean (± SEM) debiased rule cue category variance, the statistic used to measure task rule information in our previous publication from this dataset (11). As expected, results under this statistic look quite different and appear to reflect both task cue categoricality *per se* (cf. Fig. 3D) as well as the overall information about task cues (cf. Fig. 3A). (*B*) Summary (± SEM) of debiased rule cue category variance (cf. Fig. 3B,E). (*C*) Cross-area significance matrix for rule cue category variance (cf. Fig. 3C,F). (*D*) Population mean (± SEM) debiased motion category variance (cf. Fig. 4A,D). (*E*) Summary (± SEM) of debiased motion category variance (cf. Fig. 4B,E). (*F*) Cross-area significance matrix for motion category variance (cf. Fig. 4C,F). (*G*) Population mean (± SEM) debiased color category variance (cf. Fig. 5A,D). (*H*) Summary (± SEM) of debiased color category variance (cf. Fig. 5B,E). (*I*) Cross-area significance matrix for color category variance (cf. Fig. 5C,F).

**Author contributions**
M.S. and E.K.M. designed the experiments. M.S. performed the experiments and recorded the data. C.v.N. and S.L.B. spike-sorted the data. S.L.B. curated the data, and conceived and performed the analyses. S.L.B and E.K.M. wrote the manuscript.

